# Messenger RNAs transcribed from yeast linear cytoplasmic plasmids possess unconventional 5’ and 3’ UTRs and suggest a novel mechanism of translation

**DOI:** 10.1101/325316

**Authors:** Václav Vopálenský, Michal Sýkora, Tomáš Mašek, Martin Pospíšek

**Affiliations:** Laboratory of RNA Biochemistry, Department of Genetics and Microbiology, Faculty of Science, Charles University in Prague, Vinicna 5, Prague 2, 128 44, Czech Republic

## Abstract

Linear plasmids with almost identical compact genetic organization have been found in the cytoplasm of yeast species from nine genera. We employed pGKL1,2 plasmids from *Kluyveromyces lactis* as a model to investigate the previously unstudied transcriptome of yeast cytoplasmic linear plasmids. We performed 5’ and 3’ RACE analysis of all the pGKL1,2 mRNAs and found them not 3’ polyadenylated and containing mostly uncapped 5’ poly(A) leaders that are not complementary to the plasmid DNA. The degree of 5’ capping and/or 5’ polyadenylation is specific to each gene and is controlled by the corresponding promoter regions. We refined the description of the pGKL1,2 promoters and found new alternative promoters of several genes. We also provide evidence that *K2ORF3* encodes an mRNA cap guanine-N^7^-methyltransferase and that 5’ capped pGKL1,2 transcripts contain N^7^-methylated caps. Translation of pGKL1,2 transcripts is enhanced in *lsm1Δ* and *pab1Δ* strains and is independent of eIF4E and Pab1 translation factors. We suggested a model of a primitive regulation of pGKL1,2 plasmids gene expression where degree of 5’ mRNA capping, degree of 5’ non-template polyadenylation and presence of negative regulators as PAB1 and Lsm1 play an important role. Our data also suggest a close relationship between linear plasmids and poxviruses.

## Introduction

Linear plasmids have been found in the cytoplasm of yeast species from nine genera, including *Kluyveromyces*, *Debaryomyces*, *Saccharomyces*, *Saccharomycopsis*, *Wingea*, and *Pichia* (1,2). The overall genetic organization of all yeast cytoplasmic linear plasmids is identical with that of the most studied plasmids, pGKL1 (also termed K1) and pGKL2 (also termed K2), from the yeast *Kluyveromyces lactis*, where their presence is associated with the killer phenotype (3). Killer strains contain 50 to 100 copies of each of the two linear plasmids per cell (4). The pGKL1 and pGKL2 plasmids have compact and extremely AT-rich genomes sized 8874 and 13447 bp, respectively, and carry terminal inverted repeats with proteins covalently attached to their 5’ ends (5–7).

The smaller plasmid, pGKL1, contains four open reading frames (ORFs): two of them (*K1ORF2* and *K1ORF4*) encode precursors of killer toxin subunits (8–10); *K1ORF3* is involved in an immunity phenotype in an unknown manner (11), and *K1ORF1* codes for the pGKL1 plasmid-specific DNA polymerase and a terminal protein (12,13). Eleven ORFs have been reported in the larger pGKL2 plasmid, which provides vital functions for pGKL1/pGKL2 maintenance in host cell (14). Some functions have been attributed to more than half of the proteins encoded by the ORFs carried by the pGKL2 plasmid. *K2ORF2* codes for a pGKL2-specific DNA polymerase and a terminal protein (5,15). *K2ORF3* codes for a virus-like mRNA capping enzyme. A purified K2Orf3 protein (K2Orf3p) displayed 5’- triphosphatase (TPase) and RNA-guanylyltransferase (GTase) activities; however the predicted RNA cap (guanine-N^7^)-methyltransferase (MTase) activity, which is the third crucial activity of conventional mRNA capping complexes, was not detected (16,17). *K2ORF4* encodes a putative helicase probably involved in transcription of plasmid-specific mRNAs (18), *K2ORF5* codes for a single-stranded DNA-binding protein, and *K2ORF10* encodes a protein bound to the terminal inverted repeats of both pGKL plasmids. All these proteins, with the exception of K2Orf3p and K2Orf4p, are probably involved in pGKL1/2 replication (18–22). *K2ORF6* and *K2ORF7* code for putative subunits of a pGKL1/2-specific RNA polymerase (23,24). No function has been assigned to the four remaining ORFs (*K2ORF1, 8, 9*, and *11*) (Fig. 1). All ORFs are likely expressed independently. Each gene is preceded by an upstream conserved sequence (UCS) motif, which is located approximately −30 nucleotides from the putative start codon for pGKL1 UCSs (AT^A^/_C_TGA) or up to −100 nucleotides in the case of pGKL2 UCSs (ATNTGA) (for review see (2,18)). Sequences located between an AUG initiation codon and its respective UCS (inclusive) act as promoters in *sensu lato* (25–27). However, detailed mapping of the promoters recognized by the plasmid-specific RNA polymerase has not been performed.

**Figure 1.**
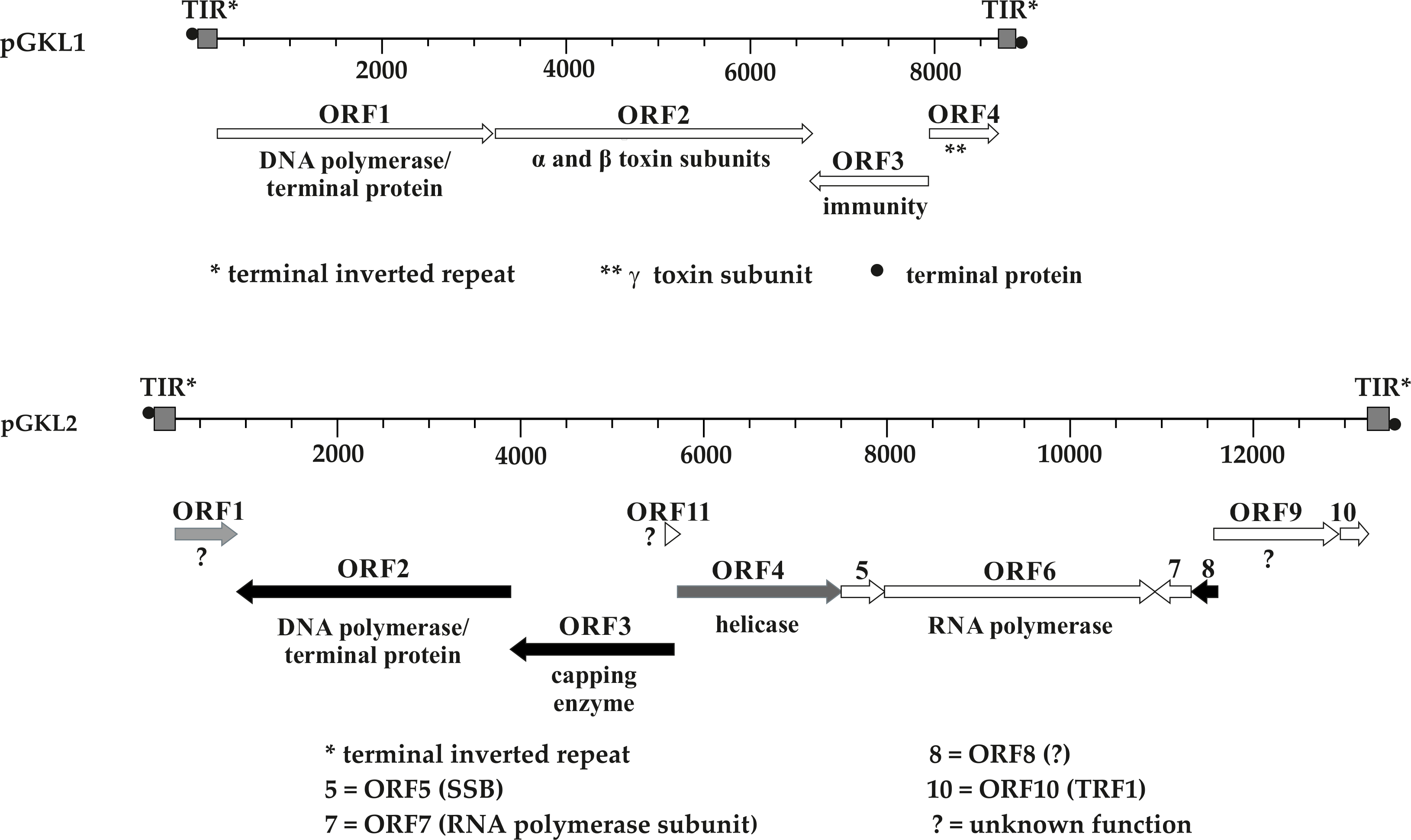
Genetic organization of pGKL plasmids. Known gene functions are indicated. Shades of gray indicate an extent of transcript capping, from black ORFs, with almost all capped transcripts, to white ORFs, with mostly uncapped transcripts. TIR - terminal inverted repeat; • - terminal protein; SSB - Single-Stranded Binding protein; TRF1 - Terminal Recognition Factor 1; ? - unknown protein function.

In summary, the yeast linear pGKL plasmids represent a model and the most studied example of a unique world of the yeast cytoplasmically localized linear double-stranded DNA plasmids. They code for their own replication and transcription and perhaps also an RNA-modification apparatus, making them remarkably independent of their host cells and evoking thoughts about their similarity to other cytoplasmically localized linear DNA genomes such as viruses belonging to the families *Poxviridae* or *Asfarviridae*. These plasmids are therefore also called dsDNA virus-like elements (VLEs) (22).

The mechanism of action of the *K. lactis* heterotrimeric killer toxin is well understood. A γ-subunit of the toxin inhibits the growth of sensitive yeast cells by cleaving their tRNAs at the 3’ side of the modified wobble nucleoside 5-methoxycarbonylmethyl-2-thiouridine (mcm5s2U), which leads to the cell cycle arrest in G1 phase (28–32). Readers are referred to references by Kast et al. and Meineke et al. for more information concerning action of the *K. lactis* toxin and related ribotoxins (31,33).

The translation of eukaryotic mRNAs is almost exclusively initiated by the binding of translation initiation factors to the 5’ cap structure. Conventional m^7^G mRNA cap formation requires three principal enzymatically catalyzed steps. During the first step, pyrophosphate is removed from the triphosphate end of the messenger RNA by TPase, GTase then transfers GMP from a GTP molecule to the new diphosphate end, and a methyl moiety is finally transferred from S-adenosyl-L-methionine (SAM) by MTase to the N^7^ position of the terminal guanosine. Conventional m^7^G-cap synthesis has been reviewed in (34,35). Whereas enzymatic activities required for canonical m^7^G-cap synthesis are highly conserved among eukaryotes and viruses, the organization of the capping apparatus itself is not conserved (36). The structural and domain organization of the capping apparatus is very diverse, particularly in viruses (37).

In this work, we focus on the yet uncharacterized transcriptome of the yeast cytoplasmic linear dsDNA plasmids and the possible mechanism of translation initiation used by the plasmid-specific mRNAs. We demonstrate that pGKL mRNAs are not 3’ polyadenylated and that most of them even lack the m^7^G cap at their 5’ ends. Surprisingly, mRNAs transcribed from pGKL plasmids contain poly(A) leaders of variable lengths at their 5’ ends that are not complementary to the plasmid DNA sequence and must thus be synthesized in a template-independent manner. We show that plasmid promoters directly determine the formation of the mRNA 5’ end leaders and their capping, pointing to a possible role of the plasmid RNA polymerase in this unusual phenomenon. We also provide the first evidence that MTase is an essential part of the K2Orf3p enzyme and that capped pGKL mRNAs are methylated at the N^7^ position of the terminal guanosine moiety. Using *in vitro* and *in vivo* approaches, we demonstrate that translation of plasmid-specific mRNAs is independent of major cellular cap-binding proteins, translation initiation factor 4E (eIF4E), and the Pab1 and Lsm1 proteins that were speculated to be possible translational enhancers and/or stabilizers of cellular and viral mRNAs bearing 5’ poly(A) stretches (38,39). Our results thus suggest the existence of a novel translation initiation mechanism.

## Materials and Methods

### General Strategy for the Modification of pGKL Plasmids Using Homologous Recombination *In Vivo*

*K. lactis* IFO1267 was transformed with a PCR-generated fragment consisting of 5’ and 3’ ends homologous to the part of the pGKL plasmid to be modified, a non-homologous part that introduces mutation/s into a plasmid-specific protein coding sequence, and a gene encoding a resistance marker (typically against G418 or hygromycin B) whose transcription is driven by the UCR of *K1ORF1* or *K1ORF2*. This type of construct was prepared by fusion PCR; for the organization of such a construct, see Fig. 2A. After transformation, yeast cells were plated onto selective agar media, and their DNA content was analyzed for the presence of the modified pGKL plasmid using agarose electrophoresis. Both modified and wild-type target plasmids were usually detected after transformation. Colonies containing both the modified and wild-type target plasmids were selected and cultivated under selective conditions for approximately 60 generations and analyzed using agarose electrophoresis. For subsequent analysis, only colonies containing a modified version of the target plasmid were used. All constructs were verified using colony PCRs and subsequent sequencing of amplified products.

**Figure 2.**
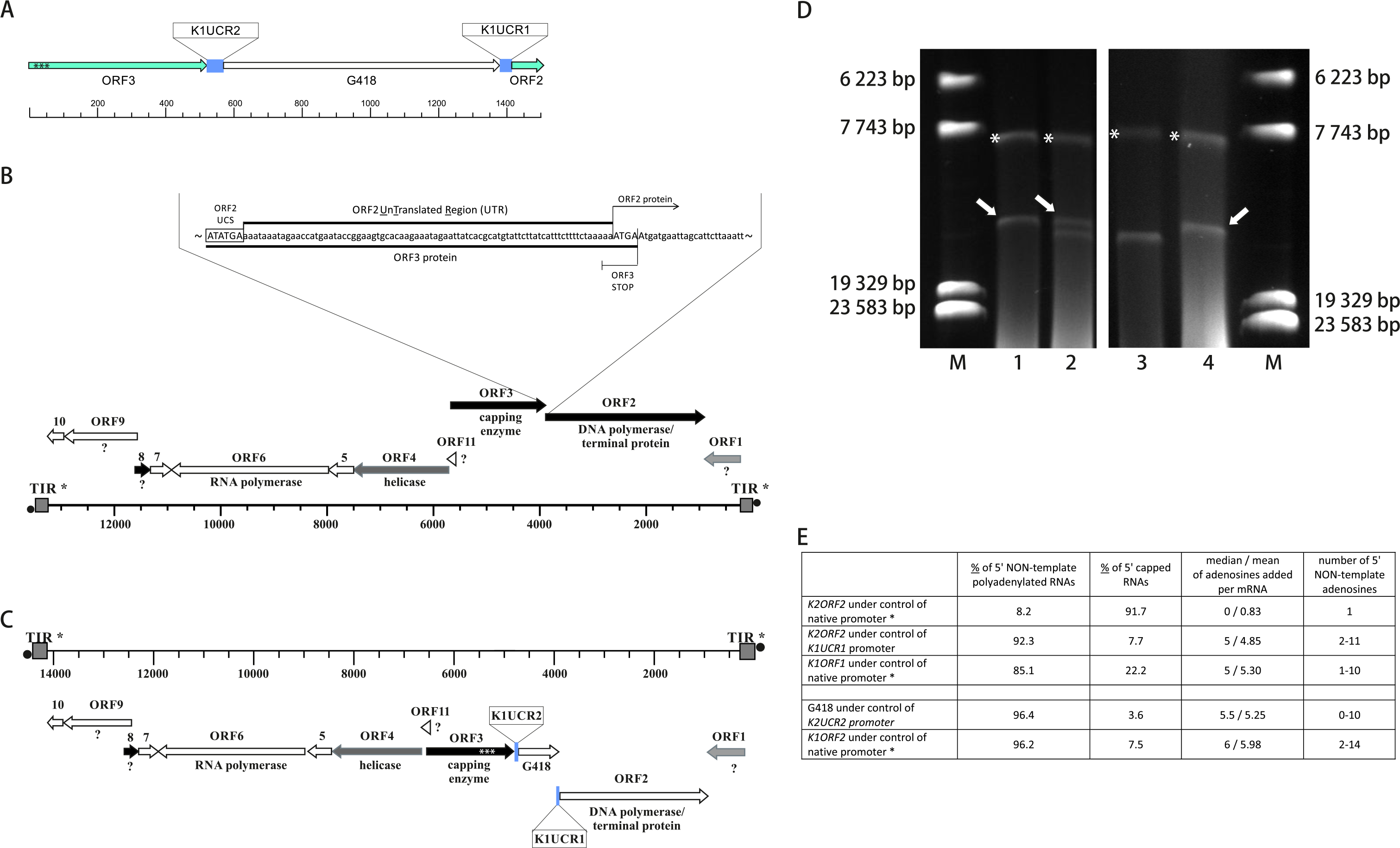
Precise manipulation of pGKL plasmids *in vivo* revealed the indispensability of the K2Orf3p MTase active site for plasmid maintenance and an essential role of pGKL promoters in mRNA capping and non-template-based 5’ polyadenylation. (A) General description of the experiment. PCR cassette used for the manipulation of pGKL2 by homologous recombination *in vivo* consists of regions homologous to the native pGKL2 plasmid (in light green), an antibiotic resistance gene (G418) under the control of the *ORF2* promoter from pGKL1 (*K1UCR2*) and the *ORF1* promoter from pGKL1 (*K1UCR1*), which will artificially control the expression of *K2ORF2*. Promoters (UCRs) are in blue, and the codons D432, G438 and D440 from the MTase active site that were mutated to Ala codons are indicated by black asterisks. (B) A closer view of the native pGKL2 region subjected to homologous recombination shows a tightly packed plasmid genome. The 3’ end of the *K2ORF3* coding region overlaps the *K2ORF2* promoter, 5’ UTR and first 4 nucleotides of the *K2ORF2* coding region. Please note that the pGKL2 plasmids displayed in panels B and C are in the reverse orientation of those in Fig. 1. Shades of gray indicate the level of transcript capping, as in Fig. 1. (C) The resulting plasmid, pRKL2-2, contains a version of *K2ORF3* with mutations in the coding region of the MTase active site (white asterisks) and two genes, aminoglycoside 3’- phosphotransferase (coding for G418 resistance) and *K2ORF2*, that are artificially controlled by the pGKL1 promoters *K1UCR2* and *K1UCR1*, respectively. Shades of gray indicate the level of transcript capping, as in Fig. 1. (D) Evidence for the vital role of K2Orf3p MTase in pGKL plasmid maintenance in cells. M - Lambda DNA/Eco130I (StyI) Marker (Fermentas); Lanes 1 and 4 - native pGKL plasmids from *K. lactis* IFO1267 (pGKL1 (8874 bp) is labeled with an asterisk, and pGKL2 (13447 bp) is labeled with an open arrow). Lane 2 - linear plasmids purified from *K. lactis* IFO1267 carrying the recombinant pRKL2-2 plasmid (14353 bp) bearing a mutated MTase active site after its cultivation for ≈ 400 generations in selective medium containing G418; Lane 3 - linear plasmids purified from *K. lactis* IFO1267 containing the pRKL2-1 plasmid (14353 bp) after cultivation for ≈ 60 generations in selective medium containing G418. Mutations in the coding region of the MTase active site spontaneously reverted to the wild-type in pRKL2-1, leading to rapid loss of the natural pGKL2 plasmid under selective conditions. Analysis of 5’ ends of pRKL2-2 plasmids revealed that non-template adenosine addition and mRNA cap synthesis are directed by linear plasmid promoters. *K2ORF2* controlled by the *K1UCR1* promoter produces fewer capped transcripts with a higher level of non-template 5’ polyadenylation, similar to wild-type *K1ORF1* transcripts and in contrast with *K2ORF2* transcripts controlled by the natural promoter. The *K1UCR2* promoter caused a similar degree of mRNA capping and 5’ non-template polyadenylation when controlling either its own gene (*K1ORF2*) or a heterologous bacterial gene coding for aminoglycoside 3’-phosphotransferase (G418 resistance). Thirteen to twenty-six independent cDNA clones were analyzed in each experiment.

### RNA Purification, Electrophoresis, Reverse Transcription, and 5’ and 3’ RACE

Total yeast RNA was purified by the hot acidic phenol procedure (40). Remaining DNA was removed by a DNA-free Kit (Ambion) according to the manufacturer’s protocol. The quality of RNA was assessed by electrophoresis according to the protocol by Masek et al. (41). For 5’ RACE, RT-PCR was carried out as follows: 0.5 μg of total yeast RNA and 0.15 μg of random primers (Invitrogen) were used for cDNA synthesis using 100 U of SSC III Reverse Transcriptase (Invitrogen) in a 20 μl reaction (50°C for 99 min). After the cDNA purification using the High Pure PCR Product Purification Kit (Roche), 800 U of recombinant TdT (Roche) and 0.5 mM dGTP (Fermentas) were used for cDNA tailing in 1x TdT buffer without CoCl_2_ for 30 min at 37°C with subsequent TdT inactivation at 70°C for 10 min. For amplification of cDNA, 2.5 μl of the reaction mixture was used for the following PCR with the universal olig2(dC) anchor primer and an appropriate gene-specific primer. In the case of 3’ RACE, 0.5 μg of total yeast RNA was polycytidinylated using a Poly(A) Tailing Kit (Applied Biosystems) with 2 mM CTP in a total volume of 25 μl for 90 min. Reverse transcription was then performed using oligo(dG)anch2 primer as indicated above. After cDNA purification using the High Pure PCR Product Purification Kit (Roche), 2.5 μl of the purified cDNA was used for a subsequent PCR reaction with the universal primer anch2 and an appropriate gene-specific primer. In both types of RACE experiments, following PCR amplification and electrophoresis, the corresponding fragments were purified from gel using a FastBack DNA minispin kit (Renogen Biolab), cloned into a pCR4-TOPO plasmid using the TOPO strategy and sequenced using the universal T7 promoter primer and/or a T3 primer. All primers used in this study are listed in Table S2.

### eIF4E Expression and Purification

The pGEX4T2::CDC33 plasmid was transformed into the expression strain *E. coli* BL21(DE3) (Merck). A 500 ml culture was grown at 28°C in 2xTY medium containing ampicillin (100 μg/ml) until the OD_600_ reached 0.20. At this point, the culture was cooled to 15°C, cultivated for 4 hours and then induced by 0.1 mM isopropyl β-D-thiogalactopyranoside. Production of recombinant protein was carried out with shaking at 15°C for another 14-16 hours. Lysates were prepared using B-PER Protein Extraction Reagents (Thermo Scientific) according to the manufacturer’s instruction and stored at −70°C for further use.

### mRNA/eIF4E Binding Assay

On days of binding experiments, bacterial lysate containing GST-eIF4E fusion protein was thawed in an ice bath, centrifuged for 10 min at 15000 g (4°C), and GST-eIF4E was purified using GST-affinity chromatography in a batch setup. In brief, 1 ml of the lysate was incubated for 80 min with 100 μl of glutathione-Sepharose 4 Fast Flow resin (GE Healthcare) at 4°C and washed four times with at least 40 volumes of ice-cold 1x phosphate-buffered saline (PBS). After the last washing step, the glutathione Sepharose binding GST-eIF4E fusion protein was resuspended in 200 μl of buffer I (20 mM HEPES, pH 7.5; 0.1 mM EDTA, pH 8.0; 100 mM KCl; and 1 mM β-mercaptoethanol) and mixed with 3 μg of DNase I-treated total RNA purified from *K. lactis* IFO1267. Next, 2 μl of the reaction mixture was removed for subsequent analysis using RT-PCR and real-time quantitative PCR (RQ-PCR). The rest of the mixture was incubated for two hours at room temperature and washed six times (with approximately 70 volumes) with buffer I. After the last washing, the glutathione Sepharose binding GST-eIF4E fusion protein and RNA and control samples were subjected to RT-PCR. All buffers used for protein purification contained Complete (EDTA-free) Protease Inhibitor (Roche), and all buffers used for handling RNA contained RNasin Ribonuclease Inhibitor (Promega).

Two microliters of reverse transcriptase reaction (obtained from eIF4E-mRNA *in vitro* binding assay) were subjected to PCR amplification as follows (5 min at 94°C; then 35 cycles of 30 sec at 94°C; 30 sec at 50°C; 45 sec at 72°C; and finally, 4 min at 72°C) using Taq DNA polymerase (Roche) and *HGT1* and *K2ORF5* gene-specific primers (listed in Table S2).

Two and half microliters of reverse transcriptase reaction (obtained from a reaction mixture before its incubation with glutathione Sepharose binding a GST-eIF4E fusion protein; see above) was subjected to Real-Time PCR amplification (5 min at 94°C; then 55 cycles of 20 sec at 94°C, 30 sec at 50°C, and 45 sec at 72°C; and finally, 4 min at 72°C) using an iCycler iQ Real-Time PCR Detection System (BioRad) and SYBR Green Master Mix (BioRad).

### Transfer of Cytoplasmic Plasmids to Yeast Strains by Incomplete Mating

The *S. cerevisiae* strain YAT547 (*kar1*) containing pGKL plasmids (42) and isogenic *S. cerevisiae* ρ^0^ strains bearing the wild-type *CDC33* gene (CWO4ρ^0^*CDC33*wt) and its temperature-sensitive mutations (CWO4p°*cdc33-1* and CWO4ρ^0^*cdc33-42*, for exact genotypes see Table S3) were cultured separately in YPD medium at 24°C overnight. Cells were resuspended the next day in 200 μl of water, and 5 μl from each of two required strains (approximately 3×10^6^ cells) were mixed on the YPD agar plate and further incubated for 5 hours at 24°C. Cells were then scraped to liquid YPD medium, incubated at 24°C with gentle shaking overnight and plated on selective plates. The resulting strains were tested for their auxotrophic markers, mating type and for killer toxin production under both permissive and non-permissive conditions. Strains showing a temperature-sensitive phenotype (when applicable), killer toxin production, growth on SD medium lacking methionine, no visible growth on SD medium lacking uracil and a MATa mating type were used for subsequent analyses.

### pGKL Plasmid Purification and Electrophoresis

For the analysis of pGKL plasmids, a modified protocol based on that of Pospisek and Palkova (43) was used. Briefly, cells were grown for three days on a dish containing the appropriate antibiotics, transferred into a microplate well and dried for 2 hours at 45°C. After they had dried completely, cells were resuspended in 40 μl of freshly prepared TESP buffer (20 mM Tris-Cl, pH 8; 50 mM EDTA-NaOH, pH 8; 2 % SDS; and 0.5 mg/ml pronase E) and dried at 37°C overnight. The sample was completely resuspended the next day in 40 μl of 1× DNA loading buffer (Fermentas). Then, 15 μl of the sample were analyzed using agarose electrophoresis (0.5 % agarose, 1 V/cm) for at least 20 hours. After the electrophoresis, the gel was incubated in a solution containing ethidium bromide (0.5 μg/ml) and RNase A (50 μg/ml) for 3 hours.

### Mutagenesis of Predicted K2Orf3p SAM Binding Site Using Homologous Recombination *in Vivo*

A fusion PCR approach was used to assembly the desired construct for SAM mutagenesis. The first primary PCR product was amplified from a native pGKL2 plasmid using Pfu DNA polymerase (Fermentas) and the primers ORF3_SAM_del_F1 and ORF3_SAMdel_R1. The second primary PCR product was amplified using Pfu DNA polymerase (Fermentas) and the primers KL_orf6C_Flag2F and ORF3_SAMdel_R2 (see Table S2) from a modified pGKL1 plasmid whose *K1ORF2* coding sequence had been replaced with G418 (unpublished data). Both primary PCR products, which had defined overlapping ends, were synthesized as follows: 5 min at 95°C; then 30 cycles of 30 sec at 94°C, 30 sec at 56.2°C, and 90 sec at 72°C; and finally, 10 min at 72°C. Then, 1 μl of each PCR mixture was used without purification for a second PCR reaction using Pfu DNA polymerase (Fermentas), the primers ORF3_SAM_del_F1 and ORF3_SAMdel_R2 and following conditions: 5 min at 95°C; then 35 cycles of 30 sec at 94°C, 30 sec at 60.1°C, and 2 min at 72°C; and finally, 10 min at 72°C). The resulting PCR product was purified using agarose electrophoresis and transformed into *K. lactis* IFO1267. After five hours of cultivation under nonselective conditions, cells were plated onto solid medium containing G418 (250 μg/ml). Site-specific integration of the PCR cassette into a pGKL2 plasmid as well as the subsequent curing of the unmodified pGKL2 plasmid was evaluated by PCR with primers (in_Kan_rev1 and in_ORF3_forw) and by sequencing of a gel-purified PCR fragment that had been amplified with primers (in_Kan_rev1 and K2_ORF3_for_seq).

### Supplemental Experimental Procedures

Please follow the Supplemental experimental procedures for the rest of the methods including Plasmid construction, Assay of enzyme-GMP complex formation, Removal of mitochondrial DNA by ethidium bromide treatment, Transformation of yeast cells, Killer production in modified CWO4 yeast strains, Assay of killer toxin activity, Statistical analysis, Expression and purification of mRNA decapping enzymes, *In vitro* processing of mRNAs by decapping enzymes, *In vitro* synthesis of capped mRNA using the vaccinia virus mRNA capping enzyme, Deletion of *PBP1*, *PAB1* and *LSM1* genes and the full list of primers, strains and plasmids used throughout this study.

## Results and Discussion

### Purification of the mRNA Capping Enzyme (K2Orf3p) and Testing its Enzymatic Activities

The *K2ORF3* gene, carried by the pGKL2 plasmid, was proposed to code for a mRNA capping enzyme similar to that of the vaccinia virus. Even though K2Orf3p apparently contains evolutionarily conserved domains corresponding to all three principal capping enzymatic activities, TPase, GTase and MTase, only TPase and GTase activities have been experimentally verified (16,17). The enzymology of mRNA cap synthesis has been reviewed in (44). A complete N^7^-methylated mRNA cap is essential for regular translation initiation in eukaryotes (45,46). We thus attempted to detect MTase activity of the purified K2Orf3 protein *in vitro*.

We produced K2Orf3p as N-terminal His-and GST-tagged proteins using an *E. coli* T7-based system or as an N-terminal His-tagged fusion protein using a baculovirus expression system. We purified the His-and GST-tagged K2Orf3 proteins to homogeneity using appropriate affinity chromatography and tested them for GTase and MTase activities. We readily detected GTase activity of the purified recombinant K2Orf3 proteins in all cases (data not shown); however, their MTase activity was not detected (data not shown). These results are in good agreement with the previously described results of Tiggemann and co-workers (17), who had shown that K2Orf3p possesses TPase and GTase activities *in vitro* but not MTase activity. They explained this result by using a C-terminal His-tag for the heterologous production of the K2Orf3p capping enzyme that may have interfered with the C-terminal MTase domain (17). We produced the putative K2Orf3p capping enzyme N-terminally fused with either His or GST tags but also failed to detect MTase activity using several *in vitro* approaches.

One possible reason we did not detect MTase activity from K2Orf3p is that the presence of an unknown stimulatory protein is required for complete and *in vitro*-detectable MTase activity of K2Orf3p, similar to a capping enzyme encoded by the vaccinia virus. The vaccinia virus encodes its own heterodimeric capping enzyme, which is composed of D1 and D12 subunits. D1 contains the domains coding for all three principal enzymatic activities required for conventional m^7^G cap synthesis arranged in an identical order to that of the domains of K2Orf3p. The D12 subunit heterodimerizes with the MTase domain of the D1 subunit and allosterically stimulates its activity (47–52). However, the MTase activity of the D1 subunit of the vaccinia virus capping enzyme is detectable even in the absence of the stimulating protein (48).

### K2Orf3p Does not Complement Conditional Mutations in *CEG1* and *ABD1* Genes

*Saccharomyces cerevisiae* yeast strains lacking active components of the mRNA capping machinery have been successfully used for studies of heterologous capping enzymes, including those from human (53,54), mouse (54), *Candida albicans* (55,56), *Schizosaccharomyces pombe* (54,57), *Encephalitozoon cuniculi* (58), *Caenorhabditis elegans* (59,60) and vaccinia virus (49,61). Putative capping enzyme K2Orf3p belongs to the D1-like group of capping enzymes due to its high similarity to the vaccinia virus capping enzyme. We therefore decided to test the ability of K2Orf3p to complement the growth of yeast cells bearing temperature-sensitive mutations in either the cap guanylyltransferase *CEG1* (GTase, (62)) or the cap methyltransferase *ABD1* (MTase, (63)) genes. We constructed yeast plasmids pYX212::NLS-K2ORF3 and pYX213::NLS-K2ORF3, which had been designed to produce K2Orf3p fused with an SV40 nuclear localization signal under the control of either a strong constitutive triose phosphate isomerase (TPI) promoter or a tightly regulated *GAL1* promoter, respectively. These plasmids were introduced into the *S. cerevisiae* strains carrying either *ceg1*^ts^ or *abd1*^ts^ temperature-sensitive alleles (62,63). We detected no growth in any of the resulting yeast strains at the restrictive temperature (data not shown). Multiple reasons can explain such a result. K2Orf3p may need to be artificially forced to interact with the C-terminal domain of RNA polymerase II or may require another co-factor for its proper function. The former approach was used to study an N-terminal fragment of the vaccinia virus capping enzyme D1 (AAs 1-545) in yeast, where it was produced as a protein fused with part of the mouse capping enzyme Mce1 (AAs 211-597). The resulting fusion protein rescued the growth of yeast strains lacking genes coding for a TPase (*cet1Δ*) and a GTase (*ceg1Δ*) (61). Saha et al. used the latter approach to show that the methyltransferase domain of the vaccinia virus capping enzyme, which is composed of the co-expressed catalytic C-terminal part of D1 and stimulatory D12 subunits, can function in lieu of the yeast cap methyltransferase Abd1 (64). A similar approach was also used to test the N^7^ methyltransferase activities of viral capping enzymes (65,66). On the other hand, some cap methyltransferases, including Hcm1 (from human), do not require any auxiliary factors to be co-expressed to support yeast growth in lieu of the Abd1 protein (54,58,66). We should notice that the vaccinia virus D1 protein itself displays mild MTase activity (48), and we complemented the temperature-sensitive phenotype, allowing us to follow the yet unobserved complementation of a partial loss of Abd1 activity in cells. Another, and perhaps the most likely, explanation why K2Orf3p does not complement the mRNA capping machinery in yeast temperature-sensitive mutants is based on the recent observation that expression of AT-rich pGKL genes in yeast nuclei does not lead to functional products due to the frequent premature 3’ polyadenylation of the resulting mRNAs (67).

### K2Orf3p MTase Active Site Integrity Is Essential for Linear Plasmid Maintenance

We and others (17) failed in numerous attempts to determine the guanosine methyltransferase activity of the K2Orf3 protein. We failed even to complement conditional mutations in the gene of yeast *ABD1*, a nuclear cap methyltransferase, with K2Orf3p targeted to the yeast nucleus. Genomes of pGKL plasmids are extremely tightly packed (Figs. 1 and 2B, C) (18). It is unlikely that the non-functional MTase region would not be eliminated from plasmids during evolution. We therefore decided to test whether the intact putative MTase is essential for the maintenance of the pGKL plasmids in *K. lactis* cells.

We developed a unique fusion PCR strategy, followed with homologous recombination *in vivo*; this approach enabled us to precisely modify the *K2ORF3* MTase coding region directly in the cytoplasmically localized pGKL plasmids (for details of the method, see Fig. 2 A-C and Materials and Methods). Using this strategy, we modified a predicted SAM-binding site of the putative *K2ORF3* MTase domain (16) by alanine scanning mutagenesis. The homologous integration of the mutational cassette (Fig. 2A) into the pGKL2 plasmid resulted in appearance of the new larger plasmid pRKL2 (<u>R</u>ecombinant *<u>K</u>*. *<u>l</u>actis* plasmid pGKL<u>2</u>, 14353 bp) in the yeast cells. This plasmid contained mutated codons encoding alanines at the positions of three amino acids (D432, G438 and D440) conserved in a predicted K2Orf3p SAM-binding site and also included the G418 resistance gene under control of the *K1ORF2* <u>U</u>pstream <u>C</u>ontrol <u>R</u>egion [UCR; a promoter in *sensu lato* and a sequence located between the AUG initiation codon and the UCS (including) of the selected ORF] (Fig. 2C and 2D). Two smaller plasmids in the cells corresponded to the natural pGKL1 (8874 bp) and pGKL2 (13457 bp) plasmids (Fig. 2D, lane 2). Randomly selected *K. lactis* clones containing all three plasmids were passaged in a selective liquid medium containing G418 (250 μg/ml) for at least 60 generations and plated on agar medium supplemented with G418 (250 μg/ml). Purified plasmids from randomly selected subclones were analyzed by agarose gel electrophoresis. In a few cases, the natural pGKL2 plasmid was lost, and only pGKL1 and recombinant pRKL2 plasmids remained in the yeast cells (Fig. 2D, lane 3). In most cases, both pGKL2 and pRKL2, containing the mutated MTase active site, replicated stably together in *K. lactis* cells passaged under selective pressure of G418 antibiotics for more than 400 generations (Fig. 2D). Four hundred generations of the mutated plasmid being favored by selective pressure and cannot outcompete the wild-type plasmid is roughly the limit of the indispensability of any analyzed pGKL ORF in this kind of experiment (68).

We purified pRKL2 plasmids from both types of yeast clones (Fig. 2, lanes 2 and 3) and sequenced the MTase coding regions of their *K2ORF3* genes. Surprisingly, yeast clones containing pRKL2 and pGKL1 only carried mutations that reversed the K2Orf3p SAM-binding site to its wild-type sequence or contained incomplete mutational cassettes that brought the same result, leaving the SAM-binding site unmodified. We called these former plasmids pRKL2-1 (Fig. 2D, lane 3). The pRKL2-2 plasmids, which were prevalent, retained the mutations in the MTase active-site-coding region and required wild-type pGKL2 to be stably maintained in the cell.

These findings clearly indicate the indispensability of the K2Orf3p SAM-binding site for maintaining pGKL plasmids in yeast cells and are consistent with previous reports stating that the entire *K2ORF3* is essential for the maintenance of pGKL plasmids in yeast cells (16).

### Linear Plasmid Transcripts are Largely Uncapped and Contain a Non-Template 5’ Poly(A) Leader

Our findings pointing to the essentiality of the K2Orf3p MTase active site for linear plasmid maintenance in yeast immediately raised the question of whether the pGKL transcripts are capped. We analyzed the 5’ ends of the pGKL transcripts using a modified rapid amplification of 5’ cDNA ends (5’ RACE) approach. Briefly, we purified total RNA from the *K. lactis* IFO1267 strain, treated it with DNase I, synthesized cDNA using SuperScript III reverse transcriptase and random primers, G-tailed the cDNA using dGTP and terminal deoxynucleotidyl transferase (TdT) and amplified the 5’ regions corresponding to transcripts from individual ORFs by PCR with an anchored oligo(dC) primer and an appropriate gene-specific primer (see primer sequences in Table S2). Reverse transcriptases, SuperScript III including, can overcome the 5’- 5’ triphosphate bond between the 5’ m^7^G cap and the first nucleotide of nascent mRNA transcripts and show whether the mRNA is capped or not (69-71). PCR fragments were verified by digestion with restriction endonucleases and cloned into a pCR4-TOPO plasmid. From 20 to 79 independent clones were sequenced and analyzed for each ORF. We used *K. lactis ACT* transcript (EnsemblFungi Id: KLLA0_D05357g), which codes for actin and is transcribed by the RNA polymerase II from a nuclear gene as a control 5’ capped mRNA.

The 5’ RACE analysis of the pGKL transcripts revealed a unique and surprising composition of 5’ ends. Eukaryotic mRNAs generally contain an m^7^G cap structure at their 5’ ends, which is a prerequisite for their stability and efficient translation initiation (72–74). Even though the pGKL plasmids encode their own putative capping enzyme (K2Orf3p), we found that few pGKL1/2 genes give rise to the frequently 5’-capped transcripts and that most of the transcripts contain a short leader of non-template adenosine nucleotides (1–20) at their 5’ ends that could not have been continuously transcribed from the plasmids (Figs. 2E, 3A and C, 7B; Tables S1A, S5). We found only two ORFs of the fifteen encoded by the pGKL plasmids, of which at least 90% of the transcripts were 5’ capped. At least 50% of capped transcripts were found in mRNAs corresponding to five ORFs, which is one-third of all ORFs encoded by the pGKL plasmids (Figs. 1, 3C; Table S1A). Non-template adenosine nucleotides were less likely added at the 5’ ends of mRNAs transcribed from ORFs whose more than 50% transcripts were capped. Conversely, longer 5’ poly(A) leaders were found in mRNAs transcribed from ORFs that yielded less frequently capped or completely uncapped transcripts (Fig. 3C; Table S1A). However, both non-templately added adenosine nucleotides and mRNA caps were frequently found within identical transcripts (Fig. 3A). We calculated the number of non-template adenosine nucleotides in the 5’ leaders of the mRNAs of each pGKL ORF as well as the minimal number of template and non-template adenosines found at the 5’ ends of their uncapped transcripts and the fractions of capped mRNAs and mRNAs containing nontemplate 5’ adenosine nucleotides. We also calculated the medians and means of the numbers of non-template adenosine nucleotides added per mRNA molecule from each pGKL ORF. These data are summarized in Fig. 3C.

**Figure 3.**
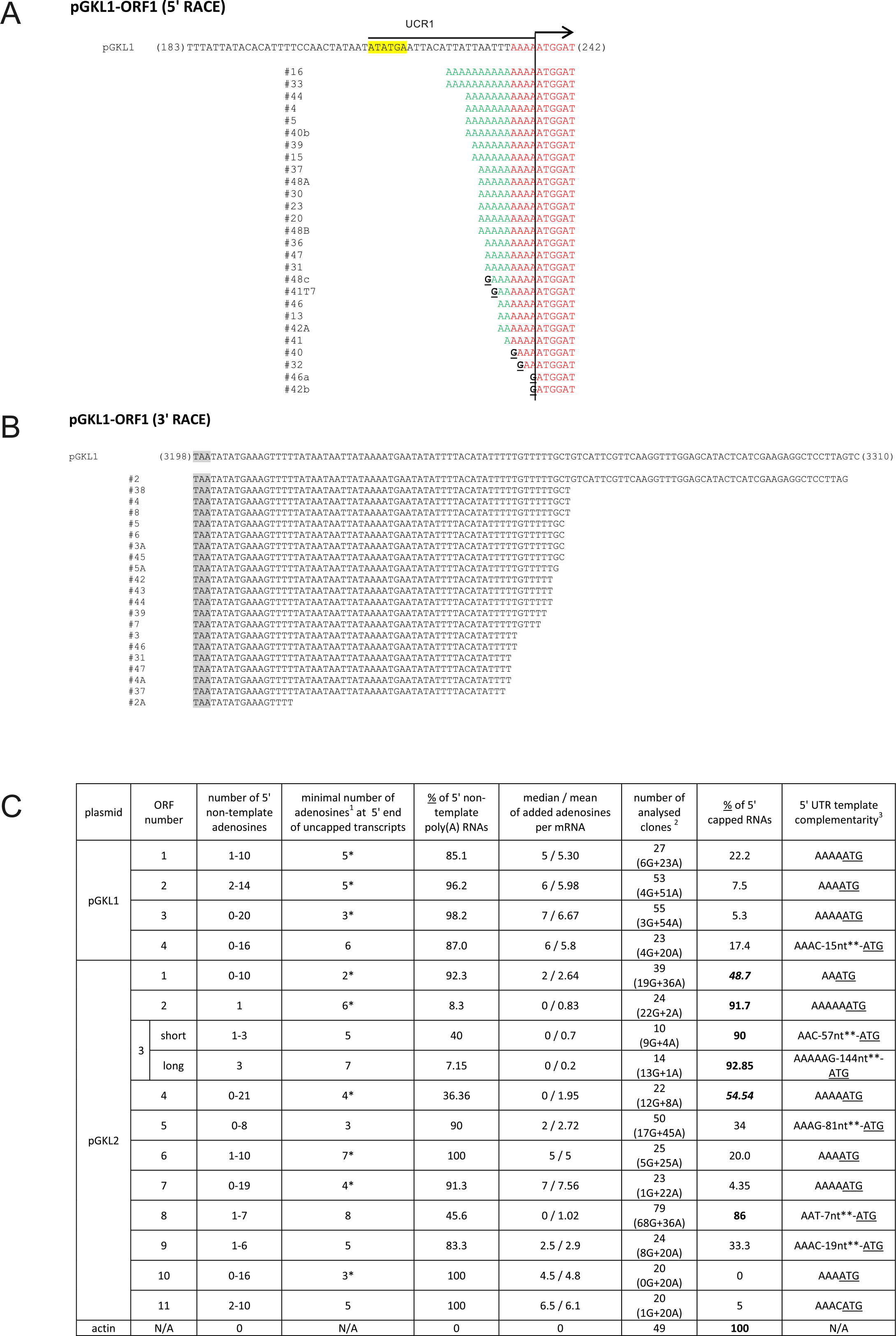
5’/3’ RACE analysis of pGKL transcripts. (A, B) Representative pictures of 5’ RACE (<u>A</u>) and 3’ RACE (<u>B</u>) analyses of individual mRNAs transcribed from *K1ORF1* encoded by the pGKL1 plasmid. The top sequence corresponds to the template (plasmid) DNA; the cDNA sequences below represent individual clones obtained by the reverse transcription of individual mRNA molecules. In the case of 5’ RACE (<u>A</u>), sequences homologous to the template (plasmid) DNA are highlighted in red, and non-template 5’ leaders of the mRNA transcripts are in green. Guanosine nucleotides corresponding to the 5’ mRNA caps are depicted in black and underlined. *K1UCR1*, which was also used as a promoter sequence in other experiments, is underlined. *K1UCS1* is highlighted in yellow. (<u>B</u>) Sequences obtained by 3’ RACE. Stop codons are highlighted in gray. For details concerning transcripts of other pGKL ORFs, please see Table S1A and the description therein. (C) Tabular summary of 5’ RACE experiments. A graphical representation of each experiment is in Table S1A and B. Each ORF is characterized by the following data: 1/ ORF number; 2/ number of 5’ non-template adenosine nucleotides found in the corresponding mRNAs; 3/ minimal number of adenosine nucleotides (calculated for both template and non-template adenosines) found at 5’ ends of uncapped transcripts; 4/ percentage of all transcripts containing 5’ non-template adenosine nucleotides; 5/ the average number (shown as the mean and median) of non-template adenosine nucleotides per mRNA molecule; 6/ number of sequenced clones used for analyses; the number of capped and 5’ polyadenylated mRNAs are depicted below the total number; 7/ percentage of all transcripts containing 5’ mRNA cap structure; 8/ part of the 5’ UTR complementary to the relevant plasmid genome. Please note that columns 4 and 7 may not sum to one hundred percent because some transcripts contain both 5’ non-template adenosines and a cap structure. ^1^ template and non-template; ^2^ G = capped transcripts; A - 5’ polyadenylated transcripts; ^3^ first templated nucleotides transcribed to the mRNA, non-template adenosines were removed; * these adenosines immediately precede the ATG sequence; ** for exact sequences, see Table S1A; transcripts from *K. lactis ACT* gene, coding for actin, were used as an internal control.

More than half of all pGKL ORFs (9) contained 5’ untranslated regions (UTRs) composed exclusively of adenosine nucleotides (Figs. 3C; Tables S1A,S5) regardless of their template or non-template origin, thus resembling adenosine-rich yeast internal ribosome entry sites (IRESs) (38). The transcripts arising from pGKL1 promoters tended to have longer and more uncapped 5’ poly(A) leaders than pGKL2 transcripts (Fig. 3C; Table S1A). Additionally, the 5’ cap is more frequently present on the transcripts containing few or zero non-template adenosine nucleotides at their 5’ ends (Table S1A). The 5’ ends of plasmid mRNAs resemble the organization of intermediate and late vaccinia virus mRNAs (75,76), late cowpox virus mRNAs (77) and even some plant mRNAs (78) that also contain non-templated poly(A) leaders at their 5’ ends. However, all these mRNA transcripts are fully capped. Considering the aforementioned examples and combining our findings with the knowledge that the domain organization of the K2Orf3p putative capping enzyme resembles that of the poxviral capping enzyme (16) allow us to speculate on the similarity between pGKL plasmids and poxviruses, at least at the level of transcription and posttranscriptional modification.

Our 5’ RACE analyses also identified the three new functional UCS sites and the corresponding transcription start sites of *K1ORF4, K2ORF3* and *K2ORF4* (Table S1A, marked as blue boxes).

### Capped pGKL Transcripts Contain an N^7^-Methylguanosine Moiety

Recent advances in the understanding of mRNA quality control and the function of closely associated cellular mRNA decapping enzymes allowed us to use *S. pombe* Rai1 and human Dcp2 proteins to directly investigate predicted N^7^-methylation at the 5’ cap guanosine of pGKL mRNAs. Expression plasmids for the production of His-tagged Dcp2 and Rai1 proteins were obtained from Prof. Mike Kiledjian (Rutgers University) and Prof. Liang Tong (Columbia University), respectively. Both proteins were produced in the *E. coli* BL21(DE3) strain and purified to homogeneity exactly as described by Xiang S. et al. in 2009 (79). We purified total RNA from the killer *K. lactis* IFO1267 strain for cap-methylation analysis. The total RNA was DNase I-treated and subsequently incubated with either purified Rai1 decapping enzyme, which removes unmethylated caps only (80), or purified hDcp2 decapping enzyme, which removes both methylated and unmethylated caps from the 5’ ends of mRNAs (81,82). To control for the proper function of both mRNA decapping enzymes, we used *in vitro*-transcribed RNA corresponding to the first 579 nt (including 32 nt of the 5’ UTR) of the *K. lactis ACT* gene, which encodes actin, (EnsemblFungi Id: KLLA0_D05357g) that had been capped using the vaccinia virus capping enzyme in either the presence or absence of SAM and subsequently incubated with Rai1 or hDcp2 decapping enzymes. The workflow of the experiment is illustrated in Fig. 4A. Treatment of *K. lactis* total RNA with hDcp2 significantly decreased the number of capped pGKL mRNAs represented by the originally highly capped transcripts from *K2ORF1* and *K2ORF2* genes, whereas treatment with Rai1 had no such effect. Identical results were obtained for the mRNA coding for actin (Fig. 4B). These results provide strong evidence that capped pGKL transcripts contain regular N^7^-methylated caps, just as the endogenous nuclear transcripts represented by actin mRNA in this experiment. *In vitro*-transcribed actin mRNA that was capped but not N^7^-methylated (G-actin RNA) served as a control of Rai1 activity. Treatment of G-actin RNA by Rai1 and hDcp2 reduced the number of capped transcripts by 43 and 29%, respectively. Conversely, caps of *in vitro*-transcribed capped and methylated actin RNA (m^7^G-actin) remained completely unaffected by Rai1 treatment, whereas hDcp2 removed 50% of all the caps (Fig. 4B).

**Figure 4.**
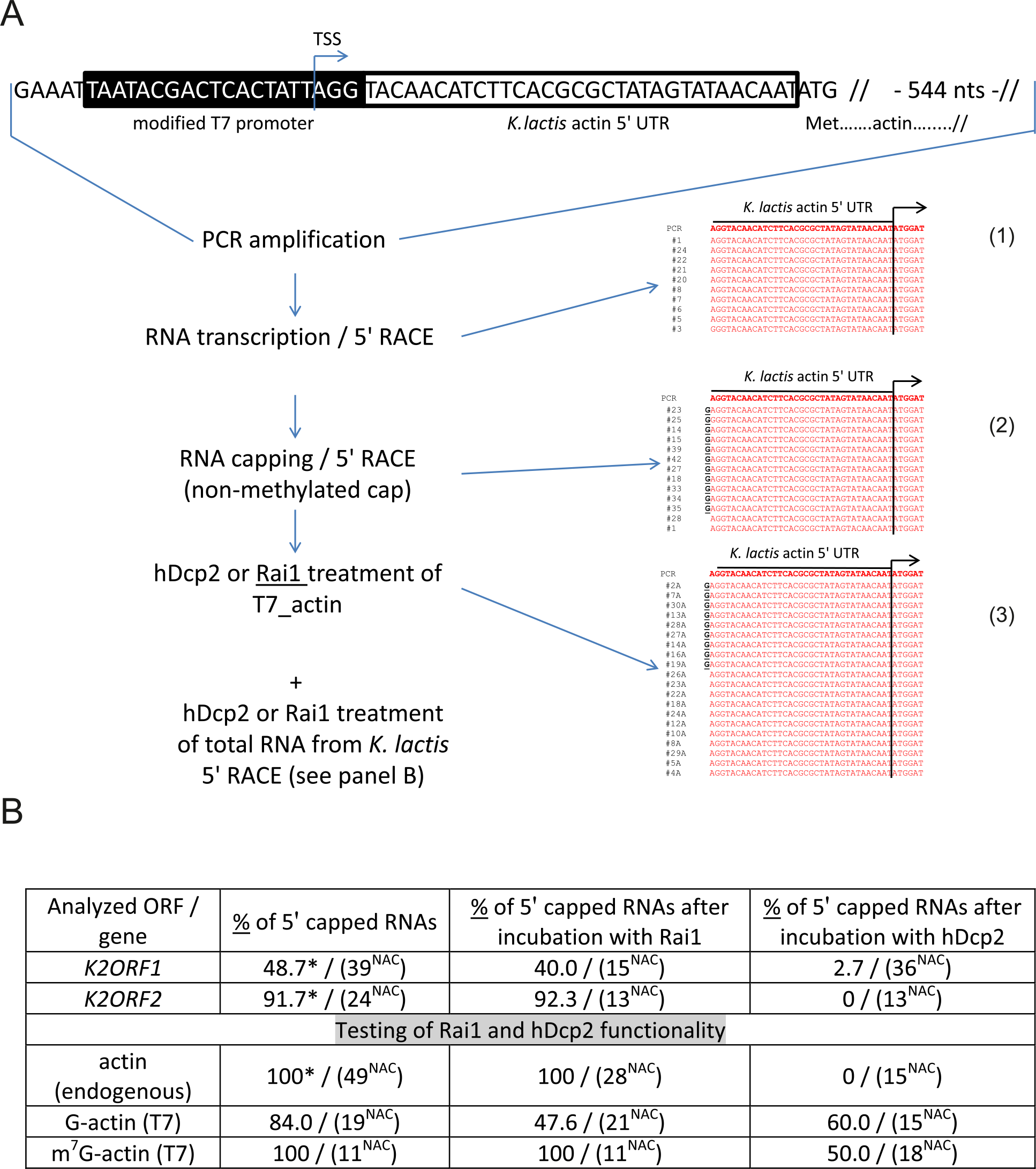
Guanosine caps at the 5’ ends of the pGKL mRNAs are N^7^-methylated. (A) Design of the control experiment. Capping of synthetic actin RNA with an unmethylated 5’ cap and 5’ RACE analysis after Rail treatment are depicted. A PCR cassette consisting of a modified T7 promoter (130) and the first 579 nt (including 32 nt of 5’ UTR) of the *K. lactis ACT* gene coding sequence was transcribed using T7 RNA polymerase. The resulting uncapped RNA molecules (1) were capped using a vaccinia virus capping enzyme in the absence (2) or presence of SAM (not shown). The RNAs were incubated with either Rail (3) or hDcp2 (not shown). The presence of either methylated (not shown) or unmethylated 5’ caps after each step was monitored by 5’ RACE followed by the cloning and sequencing of individual cDNA clones. The graphical representation of the 5’ RACE results is similar to those in Fig. 3 A. (B) Tabular representation of the 5’ RACE experiments using either total RNA prepared from *K. lactis* IFO1267 or synthetic *K. lactis* actin mRNA capped by a vaccinia virus mRNA capping enzyme in either the presence (m^7^G-actin) or absence (G-actin) of SAM. RNA preparations were analyzed untreated or treated with Rai1 or hDcp2. Numbers represent the fractions of capped RNAs from all the independent cDNA clones analyzed (numbers in brackets labeled with ^NAC^). Numbers labeled with an asterisk are replicated from Fig. 3C. NAC - number of analyzed clones.

These results also indirectly strongly support the prediction that K2Orf3p either alone or as a part of a protein complex functions as a regular capping enzyme, exhibiting all three essential enzymatic activities necessary for complete conventional m^7^G cap synthesis.

### The pGKL Promoters Determine 5’ End mRNA Capping and Non-Template Polyadenylation

Mapping of 5’ mRNA ends allows us to determine putative UCS sites, UCR regions and transcription start sites (TSSs) for all the pGKL1 and pGKL2 ORFs. *K1ORF4, K2ORF3* and *K2ORF4* contain two putative UCS sites each. We evaluated the sequences of all these UCS sites and refined their previously reported consensus sequence, ATNTGA (83), by calculating the occurrences of all 4 nucleotides in the variable N position: A = 50%, G = 11%, C = 22%, and T = 17%. The TSS (+1 nucleotide) is usually located in a short window 17 to 22 nt downstream from the 3’ end of the UCS; the median distance is 19 nt, and the extremes values are 13 and 25 nt. Transcription initiates at the adenosine nucleotide. Two exceptions were present in the minor transcripts of *K2ORF2* and *K2ORF4*; they start with G and T, respectively. The minor upstream *K2ORF4* UCS ends with a guanosine nucleotide (ATATGG), an exception to the consensus sequence, and serves as a non-canonical TSS, initiating with a T that is located only 13 nt downstream from the UCS and thus suggesting a slightly different mode of transcription initiation than in other UCRs. Most of the UTRs are very short regardless of the presence or absence of 5’ non-template leaders. Exceptions to this rule are the *K2ORF2* UTR (≈ 60 nt), *K2ORF3*_long_UTR (≈ 150 nt), *K2ORF3*_short_UTR (≈ 60 nt) and *K2ORF5* UTR (≈ 90 nt) (Table S1A). Mapping the 5’ ends of pGKL transcripts helped us to classify genes encoded by pGKL plasmids into two major groups: genes transcribed to frequently 5’-capped mRNAs with low average number of 5’ non-template adenosine nucleotides (*K2ORF2, K2ORF3, K2ORF4* and *K2ORF8*) and a larger group of genes producing transcripts less frequently 5’-capped but with a higher degree of 5’ non-template polyadenylation (Fig. 3C). UCR sequences may themselves contain signals determining the 5’ end formation of the pGKL transcripts. We took advantage of *K. lactis* strains containing altered sets of linear plasmids as described in Fig. 2. We investigated selected transcripts from the *K. lactis* IFO1267[pRKL2-1] strain (Fig. 2C and D) by 5’ RACE. The pRKL2-1 linear plasmid is a pGKL2 derivative that contains a G418 resistance marker under the control of *K1UCR2* and a *K2ORF2* artificially controlled by *K1UCR1* (Fig. 2) Indeed, we found that the presence of a cap at the 5’ end of pRKL2-1 mRNAs, as well as the presence and length of a 5’ poly(A) leader, are directly influenced by the UCR sequence used for expression. Replacement of the wild-type UCR sequence of the *K2ORF2* gene by the *K1ORF1* UCR sequence (*K1UCR1*, Fig. 3A) led to a remarkable switch from nearly fully capped and almost not 5’-polyadenylated transcripts (91.7% capped transcripts, average of added non-template adenosine nucleotides per mRNA molecule ≈ 0) to largely uncapped and 5’-polyadenylated transcripts (7.7% capped transcripts, average of added non-template adenosine nucleotides per mRNA molecule ≈ 5). These values correspond very well to the level of 5’ capping and nontemplate polyadenylation of *K1ORF1* mRNAs expressed from the pGKL1 plasmid under control of natural *K1UCR1* (Fig. 2E). The same trend was also observed for the transcripts of the resistance marker gene G418, transcription of which was controlled by *K1UCR2*. The majority of these transcripts were uncapped, but all of them were 5’ polyadenylated in exactly the same manner as natural *K1ORF2* transcripts. The same promoter had been used to drive both G418 marker gene and *K1ORF2* transcription. Additionally, the average number of nontemplate adenosine residues in the 5’ leader was comparable in both genes (G418 resistance and *K1ORF2*) controlled by *K1UCR2*. All these findings strongly support the hypothesis that differences in 5’ end formation of yeast linear plasmid mRNAs are controlled by UCR sequences and are independent of the coding sequence of a gene and its location within the pGKL plasmid system.

Our analyses of the pGKL mRNA ends and results showing the importance of the UCR region for 5’ mRNA end formation explain the surprising results of Schickel et al., who used a glucose dehydrogenase (*gdh*) reporter gene integrated into yeast cytoplasmic linear plasmids to measure the strength of several pGKL promoters. They showed that *K2UCR6* controlling the *gdh* reporter results in almost two times higher expression than when the reporter is controlled by *K2UCR10* even though these two UCRs have exactly the same length and UCS sequence (84). We found that 20% of *K2ORF6* transcripts possess a 5’ cap, whereas only uncapped mRNAs were found among *K2ORF10* transcripts (Fig. 3C, Table S1A). If a 5’ cap substantially increases the translation efficiency of its transcript, differences in cap occurrence frequency can account for differences in the translatability of pGKL plasmid mRNAs. A 5’ cap positively affects the translation of mRNAs carrying a 5’ poly(A) leader in a wheat germ extract (WGE) *in vitro* (85). Because mRNA capped with a N^7^-unmethylated cap is inefficient in translation initiation (86), a combination of our results and those of Schickel et al. (84) provides more indirect evidence for the N^7^ methylation of 5’-capped pGKL mRNAs.

### pGKL mRNAs Do Not Contain 3’ Poly(A) Tails

The unique 5’ ends of the pGKL mRNAs immediately raised a question about the organization of their 3’ ends. Previously reported indications suggested that pGKL mRNAs were polyadenylated at their 3’ ends (18,87), which is reasonable, especially in the case of non-capped pGKL mRNAs, because 3’ polyadenylation of eukaryotic mRNAs plays an important role in their stability (88,89) and the efficiency of their utilization by cellular translation apparatus (90–92). To answer this question, we performed a modified 3’ RACE where a purified and DNase I-treated total yeast RNA was 3’ tailed by an *E. coli* poly(A) polymerase using CTP instead of ATP. Following reverse transcription with an oligo(dG) anchored primer, PCR amplifications were performed with a universal anchor primer and appropriate gene-specific primers (see primer sequences in Table S2). At least twenty randomly selected 3’ RACE fragments cloned in pCR4-TOPO plasmids were sequenced for each ORF. The 3’ RACE analyses of all 15 pGKL ORFs are exemplified by the results from *K1ORF1* (Fig. 3B; Table S1C), *K2ORF5* and *K2ORF10* (Table S1C). Surprisingly, we found that pGKL mRNAs do not contain a 3’ poly(A) tail. Transcripts produced from the pGKL plasmids contain heterogeneous 3’ ends, suggesting the existence of more than one independent putative termination signal. Transcripts corresponding to only four ORFs are uniform at their 3’ ends. This group is exemplified by *K2ORF10* (Table S1C). Four pGKL ORFs gave rise to transcripts that fell into two distinct groups with regard to their 3’ UTR lengths, suggesting two independent transcription terminators. These transcripts are exemplified by those of *K2ORF5* (Table S1C). Transcripts of the remaining seven pGKL ORFs probably utilize three or more termination sites. Our initial bioinformatic analyses failed to determine the transcription termination signals. We detected only very weak RNA stem loops (ΔG from −1.0 to −4.0) at the 3’ UTR sequences that are probably not able to induce transcription termination.

### Yeast Cap-Binding Protein eIF4E Does Not Bind pGKL mRNAs *In Vitro*

The presence of a methylguanosine cap at their 5’ end and 3’ polyadenylation are common characteristics of eukaryotic mRNAs. Absence of these structures in the pGKL transcripts should lead to their decreased stability and low translatability (90,91,93). However, this behavior clearly does not occur because pGKL genes are regularly expressed. A toxin encoded by the pGKL1 plasmid is secreted in high amounts to the culture medium ((8) and next paragraph) despite the transcripts coding for the toxin subunits having the lowest cap occurrence frequency among all of the pGKL transcripts (Fig. 3C). To investigate this phenomenon, we designed a series of *in vitro* and *in vivo* experiments in which we have demonstrated that killer toxin mRNAs and probably other mRNAs starting with uncapped 5’ poly(A) leaders are translated in a novel cap-independent manner.

Translation initiation factor 4E (eIF4E) plays a crucial role in eukaryotic translation initiation. It recognizes and binds m^7^G cap structures at the 5’ ends of the eukaryotic mRNAs and brings them to the initiating small ribosomal subunits *via* its interaction with scaffold factor eIF4G. The eIF4E protein is essential in all eukaryotes, and its deletion is also lethal in yeast (94–96). The structure and function of the cap-binding eIF4E protein from the yeast *S. cerevisiae* is very well understood. Because pGKL plasmids can be transferred and stably maintained in *S. cerevisiae* (97), we decided to test possible interaction between the *S. cerevisiae* eIF4E cap-binding protein (S.c.-eIF4E) and pGKL mRNAs *in vitro*. We produced S.c.-eIF4E as an N-terminal GST fusion protein in *E. coli* and purified it by glutathione-Sepharose affinity chromatography. To test the possible binding of pGKL-specific mRNAs to the yeast eIF4E, we incubated DNase I-treated total RNA purified from *K. lactis* IFO1267 with the purified GST-S.c.-eIF4E fusion protein bound to the glutathione Sepharose. After 3 hours of incubation at room temperature, the slurry was extensively washed six times with a great excess of buffer I at room temperature and directly used for RT-PCRs with gene-specific primers. Fig. 5 clearly shows that pGKL mRNAs, represented by mRNA from *K2ORF5*, do not bind to yeast eIF4E *in vitro*, whereas cellular mRNAs, represented by mRNA coding for a high-affinity glucose transporter (*HGT1*), do. This result is further supported by qPCR analysis that revealed comparable abundances of *K2ORF5* and *HGT1* mRNAs in the *K. lactis* IFO1267 total RNA (data not shown). Interestingly, *K2ORF5* is not one of the pGKL mRNAs with the low frequency of m^7^G cap occurrence at their 5’ ends. An explanation could be that the strength of eIF4E binding to mRNA is highly influenced by the first proximal nucleotide following the terminal m^7^G, where the association constant *K_a_* is much higher for purines than for pyrimidines and slightly higher for guanosine than for adenosine (98,99). Many similar results have been obtained by other researchers; however, cap analogs were used to determine the *K_a_* and *K_d_* values in most of these studies. Adding more nucleotides after the first proximal nucleotide substantially increased eIF4E binding to the cap while still preserving the differential influence of the first proximal nucleotide on eIF4E binding (99). A short homopolymeric stretch of 5’ adenosine nucleotides thus might destabilize eIF4E binding to the mRNA cap. However, decreased eIF4E binding to the pGKL mRNAs does not substantially affect the performance of those mRNAs in translation, which suggests a novel mechanism of translation initiation independent of eIF4E cap-binding protein.

**Figure 5.**
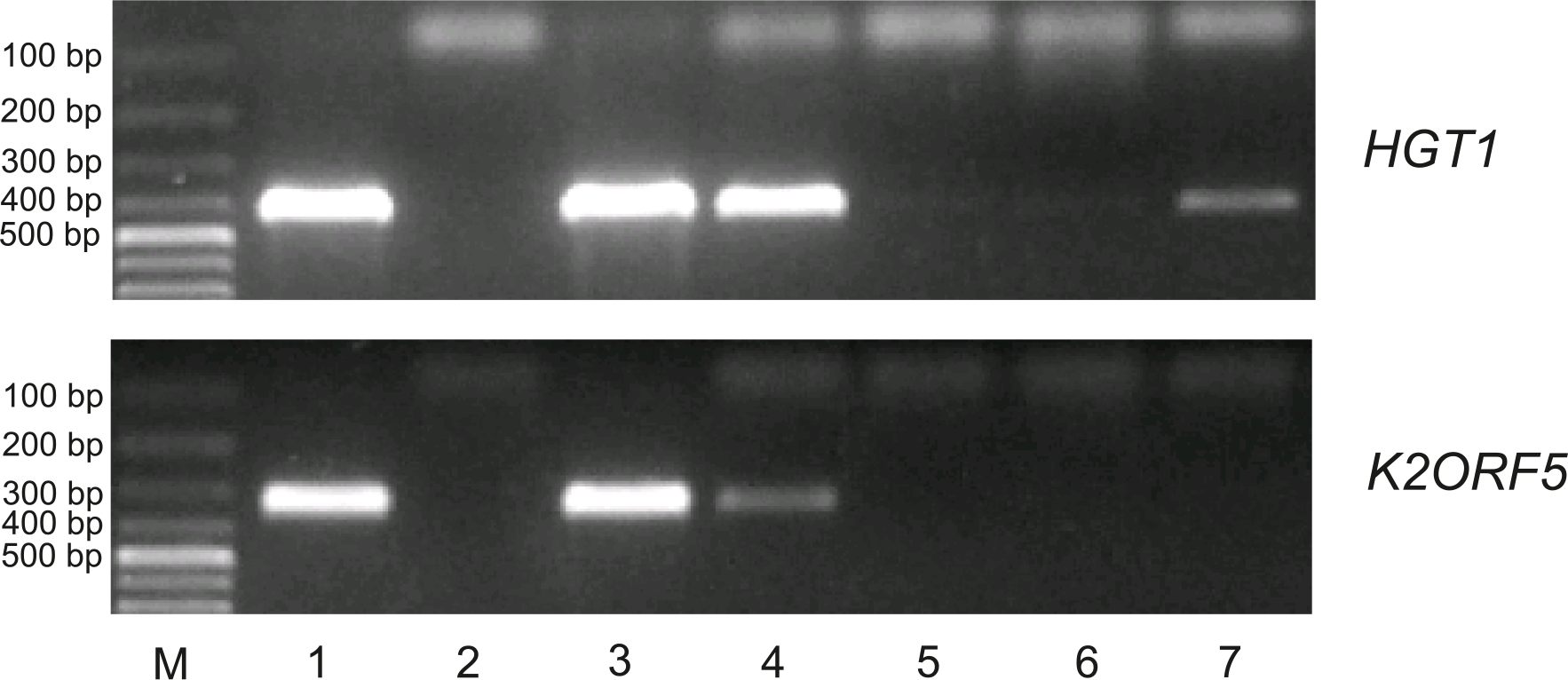
PGKL mRNAs are not bound to the yeast cap-binding protein eIF4E *in vitro*. Electrophoretic analysis of semi-quantitative RT-PCR detecting control *HGT1* mRNA (upper panel) and *K2ORF5* mRNA (lower panel) in samples as follows: Lane 1 - glutathione Sepharose binding GST-eIF4E fusion protein in the presence of *K. lactis* IFO1267 total mRNA (input); Lane 2 - same as in line 1, but reaction performed without reverse transcriptase (negative control); Lane 3 - supernatant after the first washing step (unbound mRNA); Lanes 4, 5, and 6 - supernatant after the first, third and sixth washing steps, respectively (unbound mRNA); Lane 7 - glutathione Sepharose binding complex GST-eIF4E-mRNA after the sixth washing step (bound mRNA). M - GeneRuler 100 bp DNA Ladder Plus (Thermo Scientific). PCR was performed using Taq DNA polymerase and gene-specific primers listed in Table S2.

### Killer Toxin Encoded by pGKL Plasmids is Translated by Cap-Independent Pathway

The lack of eIF4E binding to pGKL mRNAs *in vitro* prompted us to test whether these mRNAs require a functional cap-dependent pathway for their translation initiation. The reliable determination of whether a specific mRNA is translated cap-independently *in vivo* is not easy (100). We employed yeast strains bearing conditional mutations in the *CDC33* gene, which codes for eIF4E. *S. cerevisiae* CWO4 strains carrying either a *cdc33-1* or *cdc33-42* allele as a sole source of the eIF4E translation initiation factor (kindly obtained from Prof. Michael Altman) show a strong drop in protein synthesis (101,102) and a remarkable decrease in the number of polysomes (103) at an elevated temperature. We decided to transfer pGKL plasmids into these strains and determine whether a reasonable amount of the functional pGKL toxin is produced at 37°C, where Cdc33-1 and Cdc33-42 proteins are not functional and translation initiation is thus impaired (101,102). At least three genes (*K1ORF2, K1ORF3* and *K1ORF4*) encoded by the pGKL1 plasmid require sufficient expression in the yeast cells to impart killer and immunity phenotypes. The killer toxin itself is a trimeric protein comprising non-identical alpha, beta and gamma subunits that are encoded by *K1ORF2* and *K1ORF4* (Fig. 1) (8,104). Interestingly, mRNAs transcribed from all three genes belong to the group whose members have the lowest cap occurrence frequency and a high degree of 5’ polyadenylation (Fig. 3C). The presence of the killer phenotype under restrictive conditions, when cellular cap-dependent translation initiation is disabled, would thus indicate that mRNAs from at least three pGKL genes can be sufficiently and coordinately expressed in a cap-independent manner.

The artificial transfer of pGKL plasmids naturally occurring in *K. lactis* into *S. cerevisiae* is rare and difficult. In addition to other obstacles, the host mitochondria and pGKL plasmids in *S. cerevisiae* are incompatible, and the recipient strain thus must be devoid of mitochondrial DNA (ρ^0^) (42). We prepared CWO4 ρ^0^ strains solely bearing either wild-type *CDC33* or its temperature-sensitive *cdc33-1* and *cdc33-42* alleles by their long-time cultivation in the presence of ethidium bromide. To facilitate the transfer of pGKL plasmids to the new strains and to preserve their isogenic genetic background, we performed cytoduction using an incomplete mating (105) of the CWO4 ρ^0^ strains with a nuclear fusion-deficient (*kar1*) yeast strain (*S. cerevisiae* YAT547) carrying both pGKL1 and pGKL2 plasmids (42). We obtained isogenic ρ^0^ and [pGKL1/2] *S. cerevisiae* strains on the CWO4 genetic background that differed only in their *CDC33* alleles. All these strains contain both pGKL plasmids and display a killer phenotype (Fig. 6A). We required a gene coding for a yeast single-subunit protein toxin to serve as a control for cap-dependent translation. We chose a thoroughly studied K1 killer toxin encoded by the M1 viral satellite dsRNA from the yeast *S. cerevisiae*. We prepared the M1 cDNA, cloned a gene coding for the prepro-K1 killer toxin into a yeast shuttle expression plasmid (106) under the control of the strong constitutive TPI promoter (plasmid pYX212::M1) and introduced this expression plasmid into the *S. cerevisiae* CWO4 ρ^0^ strains carrying all three *CDC33* alleles described above. The resulting strains display K1 killer and immunity phenotypes at the permissive temperature (24°C) (Fig. 6B). The experiment depicted in Fig. 6 shows a comparison of the production of the *K. lactis* toxin expressed naturally from the pGKL1 plasmid and the K1 toxin expressed artificially from the nuclear plasmid pYX212::M1. The K1 toxin gene was transcribed by RNA polymerase II in the nucleus and thus provided a 5’ m^7^G-capped and 3’ polyadenylated mRNA control. We cultivated all the CWO4 ρ^0^ [pGKL1/2] and CWO4 ρ^0^ [pYX212::M1] strains in parallel to the exponential phase at the permissive temperature (24°C). All the strains were then harvested and briefly washed three times with an excess of the fresh medium prewarmed to 24°C. Half of each culture was further cultivated at 24°C. The second half was rapidly transferred to fresh medium prewarmed to 37°C and further cultivated at this elevated temperature. At this point, initial samples marked zero were taken for biomass measurements and the determination of toxin activity in the culture medium. Toxin activities were undetected in all samples at time zero and provided clear evidence that all cultures were washed sufficiently and that the toxin activities present in samples taken after 3, 6 and 12 hours of cultivation corresponded to proteins produced only in the course of the experiment (Fig. 6). The production of the *K. lactis* killer toxin in *S. cerevisiae* strains bearing pGKL plasmids was almost comparable under permissive and restrictive conditions, regardless of whether the wild-type or temperature-sensitive *CDC33* allele was present (Fig. 6A; 37°C). Production of the K1 toxin in the *cdc33-1* and *cdc33-42* yeast strains containing the pYX212::M1 plasmid vanished rapidly at 37°C, whereas the wild-type Cdc33 cap-binding protein permitted toxin production at both 24 and 37°C (Fig. 6B). The presence of the temperature-sensitive eIF4E proteins in *cdc33-1* and *cdc33-42* yeast strains was documented by their restricted growth at 37°C. Both strains containing the wild-type allele of the *CDC33* gene grew well at both 24 and 37°C (Fig. 6). The growth of CWO4 *CDC33*, ρ^0^ [pYX212::M1] was slightly slower at 24°C than at 37°C. The most likely reason is that K1 killer toxin is more active at 24°C than at 37°C, even though we cloned the coding sequence of the K1 super-killer variant that shows enhanced thermal stability (107). We previously showed that K1 preprotoxin produced artificially from the nuclear plasmid does not fully support the immunity phenotype of the host cells (106). Higher K1 toxin stability at 24°C thus probably led to the slightly slower growth. However, both cultures eventually reached comparable biomass, in contrast to the strains bearing conditional *cdc33*^ts^ alleles and cultured at a restrictive temperature (37°C), which completed the initiated cell division and ceased growth (Fig. 6B). The activity of the TPI promoter used to produce K1 toxin from the pYX212::M1 plasmid declines at the end of exponential growth (108), and this phenomenon is even more profound in the version of the TPI promoter used in the pYX212 plasmid (unpublished data). The growth of the CWO4 *CDC33*, ρ^0^ [pYX212::M1] strain is slightly faster at 37°C than at 24°C, causing the cells to reach early stationary phase in 8 hours (instead of 12 at 24°C); this difference explains why K1 toxin production in the *CDC33* wild-type strain ceases in samples grown for 12 hours at 37°C (Fig. 6B).

**Figure 6.**
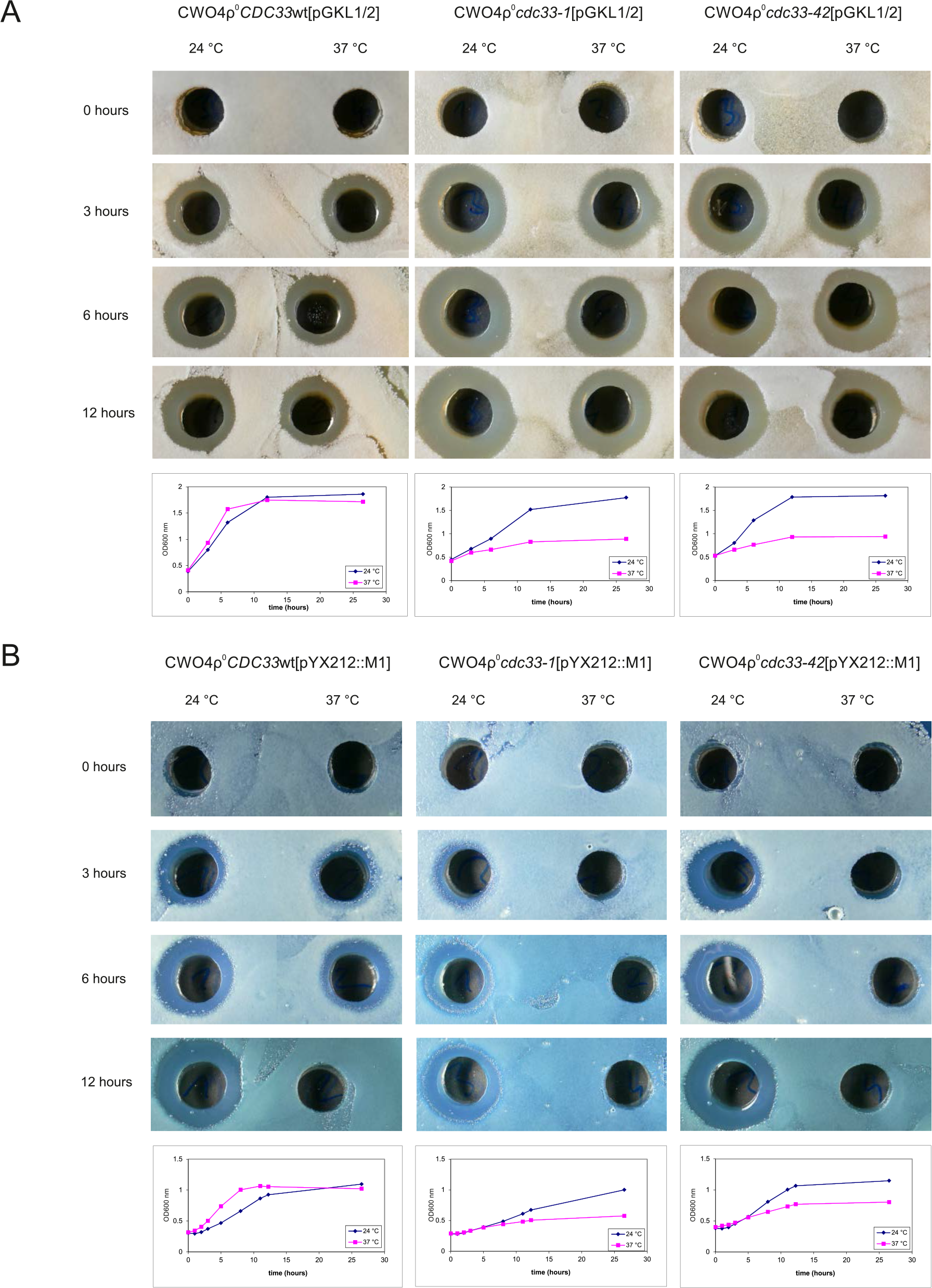
Killer toxin encoded by pGKL plasmids is translated by the cap-independent pathway. (A) Production of pGKL killer toxin in *S. cerevisiae* CWO4ρ^0^ strains bearing cytoplasmic pGKL plasmids and different *CDC33* alleles. The pGKL toxin-coding mRNAs do not contain 3’ poly(A) tails and are mostly 5’ uncapped and 5’ polyadenylated. (B) Production of K1 killer toxin in control *S. cerevisiae* CWO4ρ^0^ strains containing pYX212::M1 plasmid and different *CDC33* alleles. The control mRNA coding for K1 toxin is transcribed in the nucleus by RNA polymerase II. In both cases, the results are shown for *S. cerevisiae* CWO4ρ^0^ strains bearing either the wild-type *CDC33* (eIF4E) gene (*CDC33*wt) or temperature-sensitive mutations in the *CDC33* (eIF4E) gene (*cdc33-1* and *cdc33-42* alleles) grown under both permissive (24°C) and non-permissive (37°C) conditions. Aliquots were taken at 0, 3, 6 and 12 hours, and the culture medium was filter-sterilized, diluted 10x, and assayed for the presence of the killer toxin activity by an agar well diffusion test using a lawn of *S. cerevisiae* S6/1 sensitive strain cells. Growth curves of all strains at permissive (blue) and non-permissive (purple) temperatures are shown below each of the corresponding sets of killer toxin experiments.

The most straightforward explanation of the obtained results is that whereas in this experiment, K1 toxin mRNA is transcribed from the shuttle plasmid under the control of a TPI promoter by the nuclear RNA polymerase II and is thus 5’ capped, 3’ polyadenylated, and translated exclusively by the canonical cap-dependent pathway, pGKL1 toxin mRNAs retain their ability to be translated even when 5’ mRNA cap recognition is severely impaired. The pGKL1 mRNAs are transcribed by the pGKL plasmid-specific RNA polymerase encoded by *K2ORF6/7* and contain unusual 5’ stretches of non-template adenosine nucleotides rarely linked with the terminal m^7^G caps (Figs. 1 and 3C). Our experiments, both *in vitro* (Fig. 5) and *in vivo* (Fig. 6), strongly suggest that these 5’ terminal structures of the pGKL mRNAs support a novel manner of translation initiation independent of the active eIF4E protein. The active translation of 5’ polyadenylated mRNAs, which do not bear 5’ caps and 3’ poly(A) tails, is further supported by our observation that these mRNAs are present in polysomes in both the yeast *K. lactis* IFO1267 naturally bearing pGKL plasmids and the engineered *S. cerevisiae* CWO4 ρ^0^ [pGKL1/2] strains (data not shown).

### Pab1 and Lsm1 Deletions Support the Translation of pGKL mRNAs

Poly(A) tracts within the 5’ UTRs and the 5’ poly(A) leaders of the eukaryotic and viral mRNAs have been identified as enhancers of mRNA translation in various examples (38,85,109,110) as well as translational repressors in other cases (111,112). Some studies even show that poly(A) stretches within 5’ UTRs may serve as translational enhancers until they reach a certain threshold length. Further increases in the number of consecutive adenosine nucleotides beyond the threshold length lead to translational repression (113). Poly(A)-binding protein 1 (PABP1) plays an important role in many of these studies and biological examples. PABP1 is an abundant general translation initiation factor that binds to 3’ poly(A) tails, a common mark of most eukaryotic mRNAs (114). In addition, PABP1 serves as a specific regulatory protein and has an important role in mRNA stability and turnover. The importance of PABP1 in cellular translation and other vital processes is underlined by growing evidence that it is targeted by a wide range of viruses aiming to take over host protein synthesis or to escape the cellular antiviral defense (115–118).

To investigate the possible effects of yeast PABP1 protein (Pab1) on the expression of pGKL genes, we deleted the *PAB1* gene from *K. lactis*. Because no experimental data describing *PAB1* deletion in the yeast *K. lactis* were available, we followed a strategy used for the deletion of *PAB1* in the yeast *S. cerevisiae* (119). Deletion of *PAB1* gene is lethal in *S. cerevisiae* (94,120) but this lethality can be suppressed by the simultaneous deletion of the gene coding for Pab1p-binding protein (Pbp1) (119). We disrupted the *PBP1* gene in the chromosomal DNA of the *K. lactis* IFO1267 strain using a loxP-G418-loxP system (121) and obtained the *K. lactis* IFO1267 *pbp1*::G418 strain. The G418 cassette was removed from the genome by an artificially produced Cre-recombinase, providing the strain *K. lactis* IFO1267 *pbp1Δ*. We used the same strategy to introduce a *PAB1* deletion and to obtain the required strain, *K. lactis* IFO1267 *pbp1Δ pab1Δ*. Another protein that stabilizes some viral 5’ poly(A) tracts and thus may also affect translation of the respective mRNAs is Lsm1 (39). We disrupted the *LSM1* gene in the *K. lactis* IFO1267 genome using a strategy similar to that described for *PBP1* deletion. The *K. lactis pbp1Δ pab1Δ* double-mutant strain is viable but exhibits a slow-growth phenotype, similar to the reported behavior in *S. cerevisiae* (119).

Deletions of *LSM1* and *PBP1* from *S. cerevisiae* decrease growth rates (122,123). In contrast, we detected only minor decrease in the growth rate of the *lsm1Δ* strain and no difference in the growth rate of the *pbp1Δ* strain from the growth rate of their parental wild-type strain, *K. lactis* IFO1267 (Fig. 7A). Disruption of any of the three genes tested enhanced production of the toxin encoded by the linear cytoplasmic plasmid pGKL1, and the production was most enhanced in the *lsm1Δ* strain. The increased toxin production by the slow-growing *K. lactis* IFO1267 *pbp1Δ pab1Δ* strain was more noticeable when toxin production to the culture medium was quantified as the diameter of the inhibition zone on the lawn of the sensitive yeast strain per production cell (Figs. 7A, S1) The Lsm1 protein is part of the complex of cellular mRNA decapping activators employed by positive-strand viruses to promote their translation and replication (reviewed in (124)) and might stabilize orthopoxviral mRNAs containing 5’ poly(A) leaders (39). Despite the many similarities between poxviruses and the apparatuses of pGKL transcription and RNA modification (22), the simplest explanation for increased pGKL toxin production in the *lsm1Δ* strain might be the stabilization of pGKL mRNAs in the cell due to prominent involvement of the Lsm1 protein in a cytoplasmic deadenylation-dependent mRNA decay pathway (reviewed in (125)). We calculated the lengths of the continuous 5’ poly(A) leaders of the pGKL mRNAs while considering both non-template and template-coded adenosine nucleotides. While the maximal length of the 5’ poly(A) leaders can reach 26 nucleotides, their average and median lengths are much lower. The median lengths of 5’ poly(A) leaders in transcripts from only two genes, *K1ORF3* and *K2ORF7*, are 12 adenosine nucleotides. Transcripts from *K1ORF1, K1ORF2* and *K2ORF11* have 5’ poly(A) leaders with median lengths of 10 adenosine nucleotides. All other pGKL transcripts have shorter 5’ poly(A) leaders (Tables S1A and S5). These results agree very well with the genome-wide study of Xia et al., who showed that the presence of poly(A) stretches in 5’ UTRs in *S. cerevisiae* transcripts positively correlates with enhanced protein synthesis until the length of the 5’ poly(A) regions reaches at least 12 consecutive adenosine nucleotides (113), which is also the optimal length for the binding site of yeast Pab1 (120). Pab1 might function as a negative regulator that can, due to the documented sharp increase in its binding affinity to poly(A) stretches ranging from 8 to 12 adenosine residues in length (120), further modulate the expression of pGKL genes. Such Pab1 activity was suggested for larger set of yeast genes and can be demonstrated in mRNA coding for cyclin *PCL5*, which contains the longest yeast 5’ UTR poly(A) region and shows inefficient translation and extremely low ribosomal occupancy (113). In mammalian cells, PABP1 binds as a part of the negative autoregulatory complex to the adenosine-rich elements within the 5’ UTR of its own mRNA (111). mRNAs containing long uncapped poly(A) leaders (25 nt long) are efficiently translated in WGE lysates and, in contrast to other mRNAs, do not commonly decrease in translation rate at high mRNA concentrations (85). This behavior suggests a lower dependence of such mRNAs on translation factors. Indeed, mRNAs containing long 5’ poly(A) leaders can efficiently enter translation, even in the absence of the otherwise essential translation factors eIF4F, eIF3 and PABP1 (109). Late poxviral mRNAs containing long 5’ poly(A) leaders have a low requirement for eIF4F and might have a low requirement for PABP1 (113,126). Interestingly, pGKL1 mRNAs coding for toxin and immunity phenotypes contain a higher fraction of mRNAs with poly(A) leaders that are at least 12 residues long (Tables S1A and S5). These mRNAs, which perhaps normally have Pab1 bound to their 5’ UTRs, may become unblocked and available for translation in the *K. lactis pbp1Δ pab1Δ* strain, thus increasing overall pGKL toxin production. The control strain, *K. lactis pbp1Δ*, showed no growth defects and the lowest enhancement of pGKL toxin production Fig. 7A. Similarly to its human ortholog ataxin-2, yeast Pbp1 protein is involved in a wide range of cellular process; malfunction of ataxin-2 causes serious diseases in humans, including neurodegenerative spinocerebellar ataxia type II (127). Pbp1 (ataxin-2) functions are particularly associated with mRNA stability and translation, and Pbp1 is a negative regulator of a poly(A) nuclease (128), which might be a reason for the enhanced production of the pGKL toxin in the *pbp1Δ* strain. To exclude the possibility of changes in 5’ poly(A) leader length and m^7^G capping of pGKL transcripts in the *pbp1Δ pab1Δ* double-mutant strain, we analyzed 5’ ends from *K1ORF2* and *K1ORF3* mRNAs coding for the toxin α/β-subunit and immunity, respectively, by the 5’ RACE. As clearly seen in Fig. 7B, deletion of the *PAB1* gene does not remarkably affect the structure of the 5’ ends of the pGKL mRNAs.

**Figure 7.**
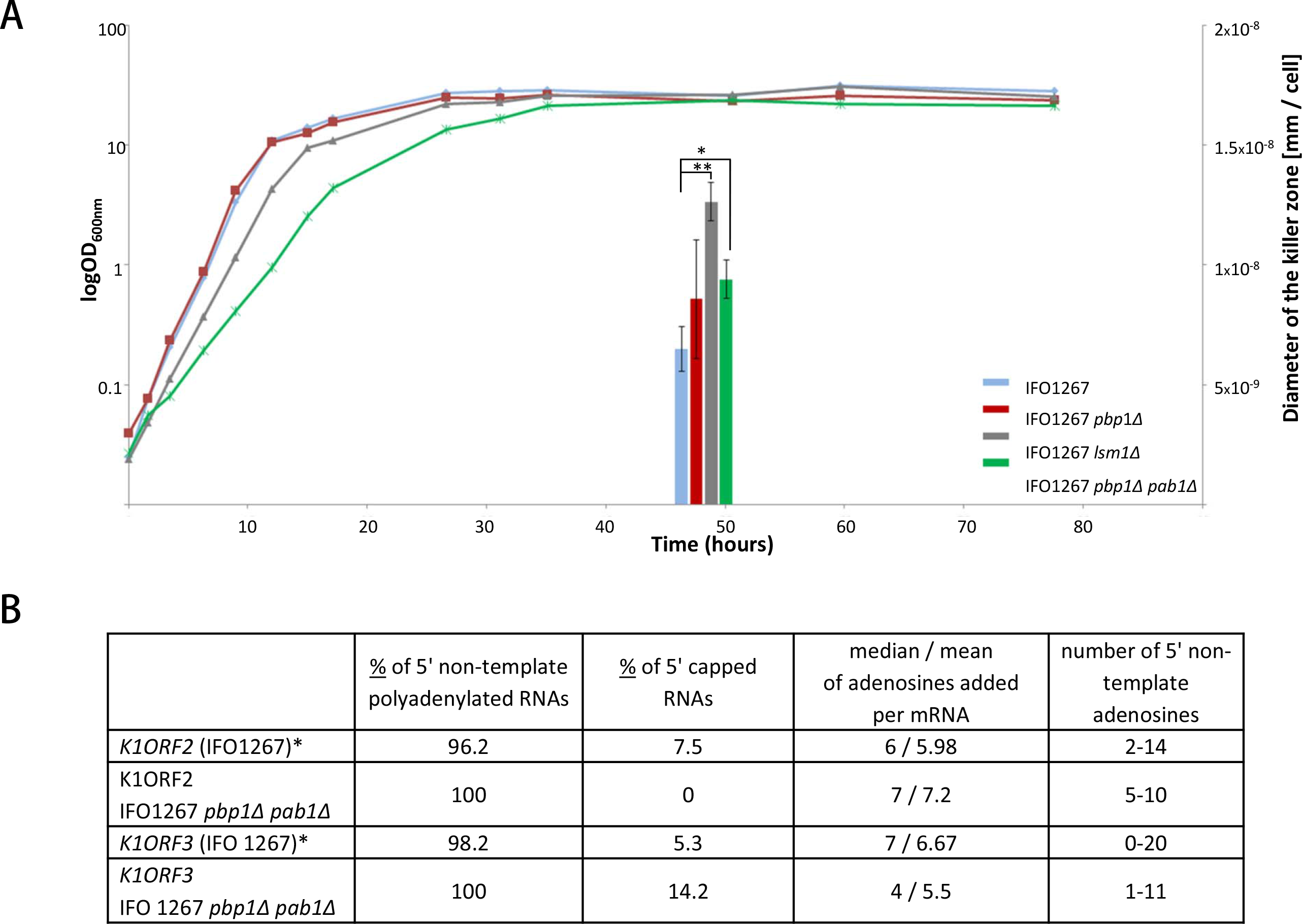
Translation of the *K. lactis* killer toxin is enhanced in *LSM1* and *PAB1* deletion strains. (A) Growth curves and production of the pGKL1 killer toxin from *K. lactis* IFO1267 (blue), IFO1267 *pbp1Δ* (red), IFO1267 *lsm1Δ* (yellow) and IFO1267 *pbp1Δ pab1Δ* (green) strains. All strains were cultivated in YPD medium at 28°C for approximately 80 hours, and their OD_600 nm_ and toxin production were monitored. Production of the pGKL1 killer toxin into culture medium was assayed by a well diffusion test on YPD agar plates with a lawn of the *S. cerevisiae* S6/1 sensitive strain (Fig. S1). Toxin levels in a culture medium in early stationary phase (~51 hours), when all the cultures reached comparable OD_600_, are depicted. Killer zones were measured using 100x diluted filter-sterilized culture medium, and the average width of the killer zone per cell was calculated for each strain, plotted with standard deviation and statistically analyzed (** p < 0.01, * p < 0.05) (B) 5’ RACE analysis of pGKL mRNAs from the wild-type and *PAB1* deletion strains. 5’ RACE analysis of *K1ORF2* and *K1ORF3* mRNAs was performed using total RNA prepared from *K. lactis* IFO1267 and its double-deletion mutant *K. lactis* IFO1267 *pbp1Δ pab1Δ*. The individual columns represent the following quantities: 1/ percentage of all transcripts containing 5’ non-template adenosine nucleotides; 2/ percentage of all transcripts containing a 5’ cap structure; 3/ the average number (shown as the mean and median) of non-template adenosine nucleotides per mRNA molecule; 4/ the number of 5’ non-template adenosine nucleotides found in the corresponding mRNAs. Data related to the wild-type strain *K. lactis* IFO1267 are replicated from Fig. 3C.

## Conclusions

We employed pGKL plasmids from the yeast *Kluyveromyces lactis* as a model to investigate and present here a complex view of the largely unknown transcriptome of yeast cytoplasmic linear plasmids. Although previous experiments reported the binding of pGKL mRNAs to an oligo(dT) column (18), we show that the pGKL transcripts are not polyadenylated at their 3’ ends; however, they frequently contain uncapped 5’ poly(A) leaders that are not complementary to the plasmid DNA. The degree of 5’ capping and/or 5’ polyadenylation is specific to each gene and is most likely controlled by short UCRs preceding the corresponding AUG start codon. This transcriptome analysis allowed us to refine the description of the pGKL promoters and revealed new alternative promoters for the *K1ORF4, K2ORF3* and *K2ORF4* genes. We also provide evidence that *K2ORF3* codes for the long sought mRNA cap guanine-N^7^ methyltransferase and that capped pGKL transcripts contain N^7^-methylated caps at their 5’ ends. Translation of pGKL transcripts is independent of eIF4E and Pab1 translation factors. With regard to yeast (113) and mammalian (111,112) mRNAs containing long poly(A) stretches in their 5’ UTRs, we suggest that Pab1 serves as a negative regulator of the pGKL mRNAs containing 5’ poly(A) leaders. It is tempting to speculate that differences in the median lengths of the 5’ mRNA poly(A) leaders of each gene (Tables S1A, S5) might reflect the degree of inhibition of the corresponding gene expression by Pab1, which might also be an evolutionary force driving the selection of the appropriate UCRs. We hypothesize that the small and extremely compact genome of pGKL plasmids uses different levels of 5’ capping and 5’ polyadenylation to finely tune the expression of each gene to proper levels to facilitate the formation of functional protein complexes and to keep pGKL gene expression in harmony with the host cell. Some initial evidence supporting this hypothesis comes from comparing our data with that of Schickel et al., as discussed earlier (84). We also provide data further supporting an evolutionary relationship between a group of the cytoplasmic yeast linear plasmids and poxviruses (22). This connection is fascinating because members of the *Poxviridae* family have been found only in vertebrates and arthropods, and some of them cause highly contagious and serious diseases of humans and animals. Parallel studies of yeast cytoplasmic linear plasmids and poxviruses may lead us to the origin of eukaryotic viruses. The pGKL plasmids could also help us better understand basic processes and host cell-virus interactions that are not easily studied in mammalian systems. Last but not least, *K. lactis* is one of the most important yeast for research and industrial biotechnology (129), and our results suggest that *lsm1Δ* strains can greatly improve the production of heterologous proteins from pGKL-based expression plasmids.

## Acknowledgements

We thank Petra Studnickova, Vlasta Pelechova and Natalie Suchankova (Charles University, Czech Republic) for their help. We thank Prof. Michael Altman (University of Bern, Switzerland), Prof. Norio Gunge (Sojo University), Prof. Mike Kiledjian (Rutgers University), Prof. Liang Tong (Columbia University), and Prof. Beate Schwer (Weill Cornell Medical College) for yeast strains and plasmids.

## Supplementary information

**Figure S1.**
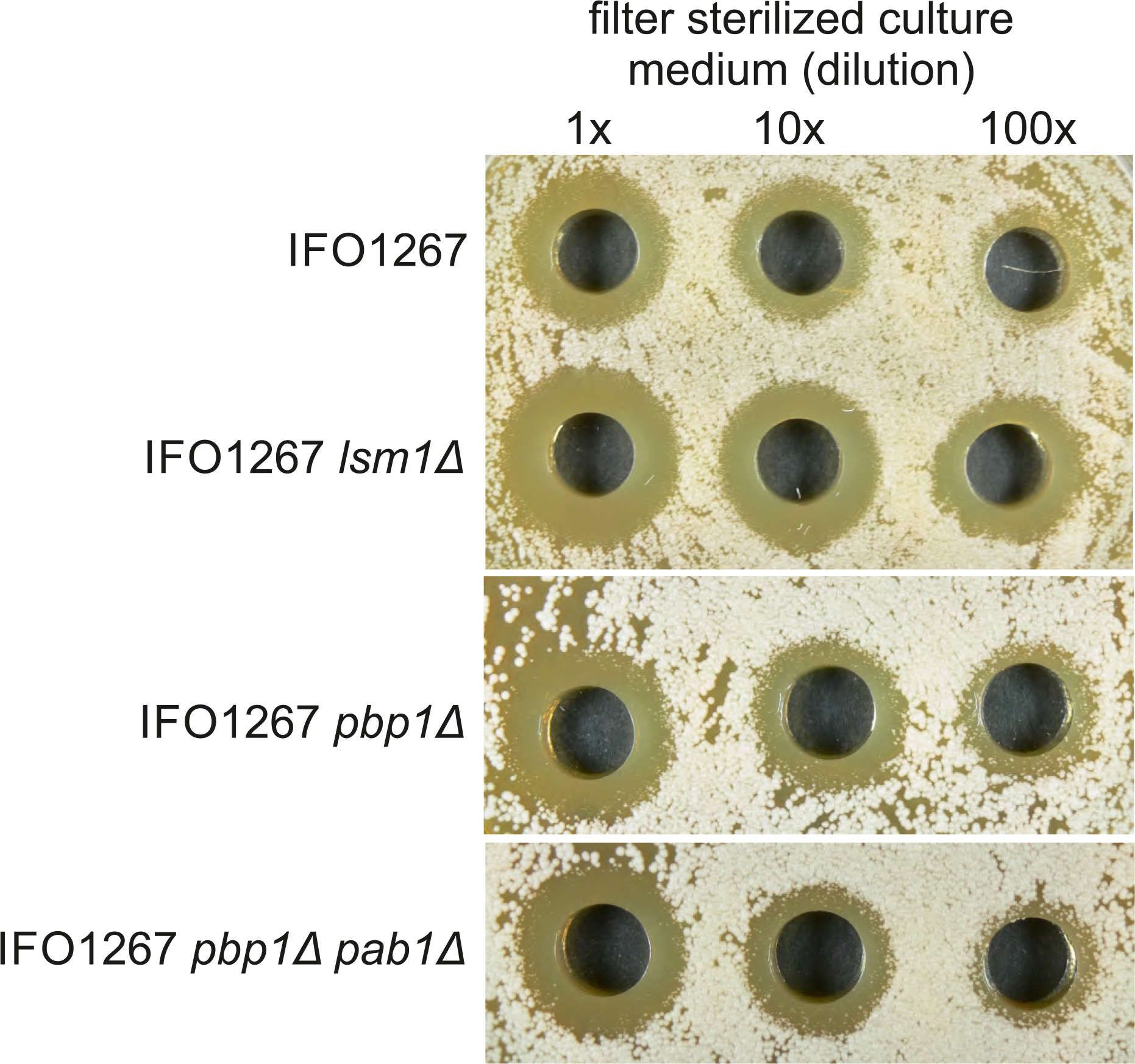
Production of the pGKL1 killer toxin in early stationary phase. Production of the pGKL1 killer toxin from *K. lactis* IFO1267, IFO1267 *pbp1Δ*, IFO1267 *Ism1Δ* and IFO1267 *pbp1Δ pab1Δ* strains to culture medium in early stationary phase (~51 hours), when all the cultures reached comparable OD_600_ (see Fig. 7). Production of the pGKL1 killer toxin into culture medium was assayed by a well diffusion test on YPD agar plates with a lawn of the *S. cerevisiae* S6/1 sensitive strain. Culture medium was used either undiluted (1x) or diluted 10x and 100x with sterile YPD medium. This figure refers to Fig. 7.

**Table S1.**
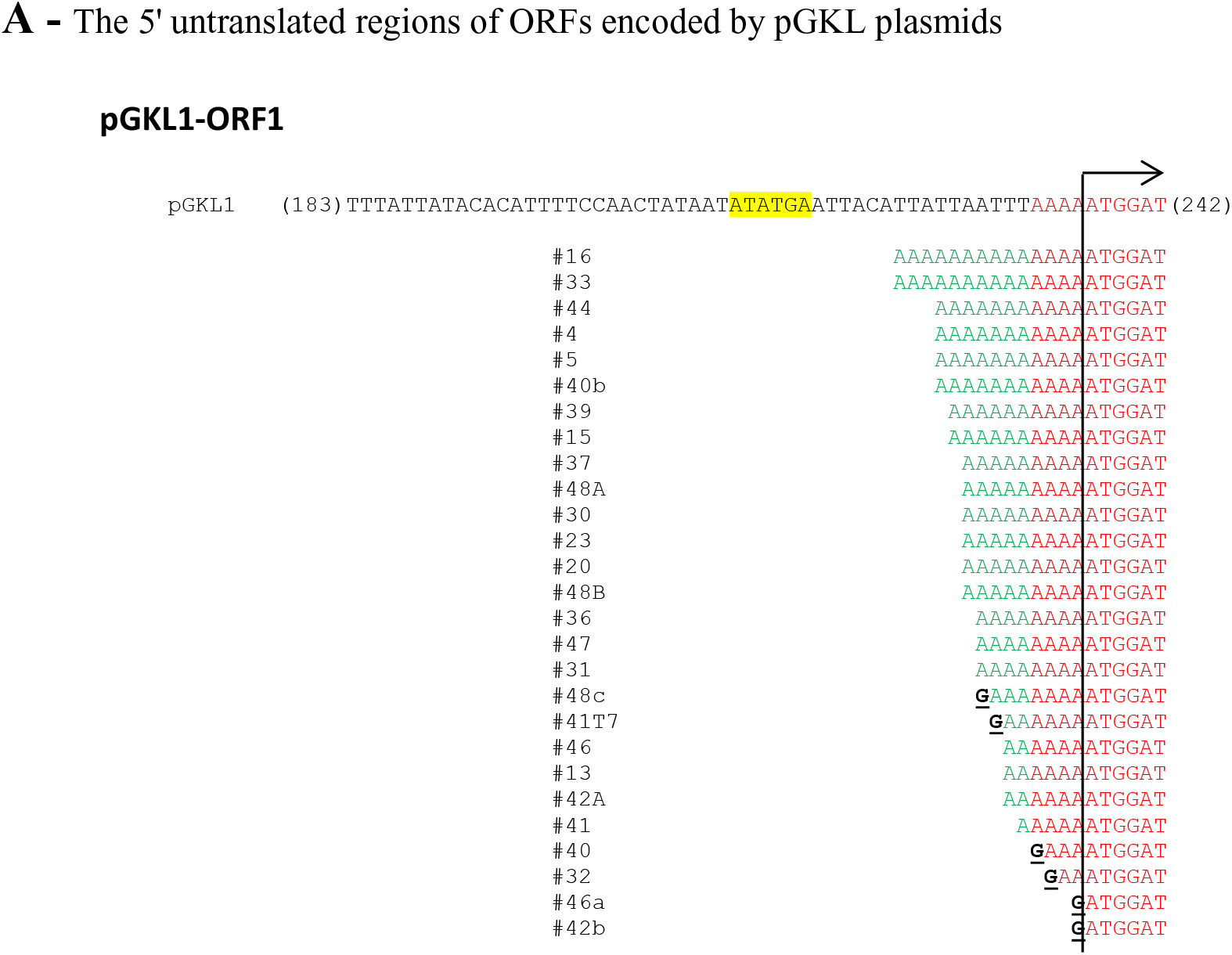
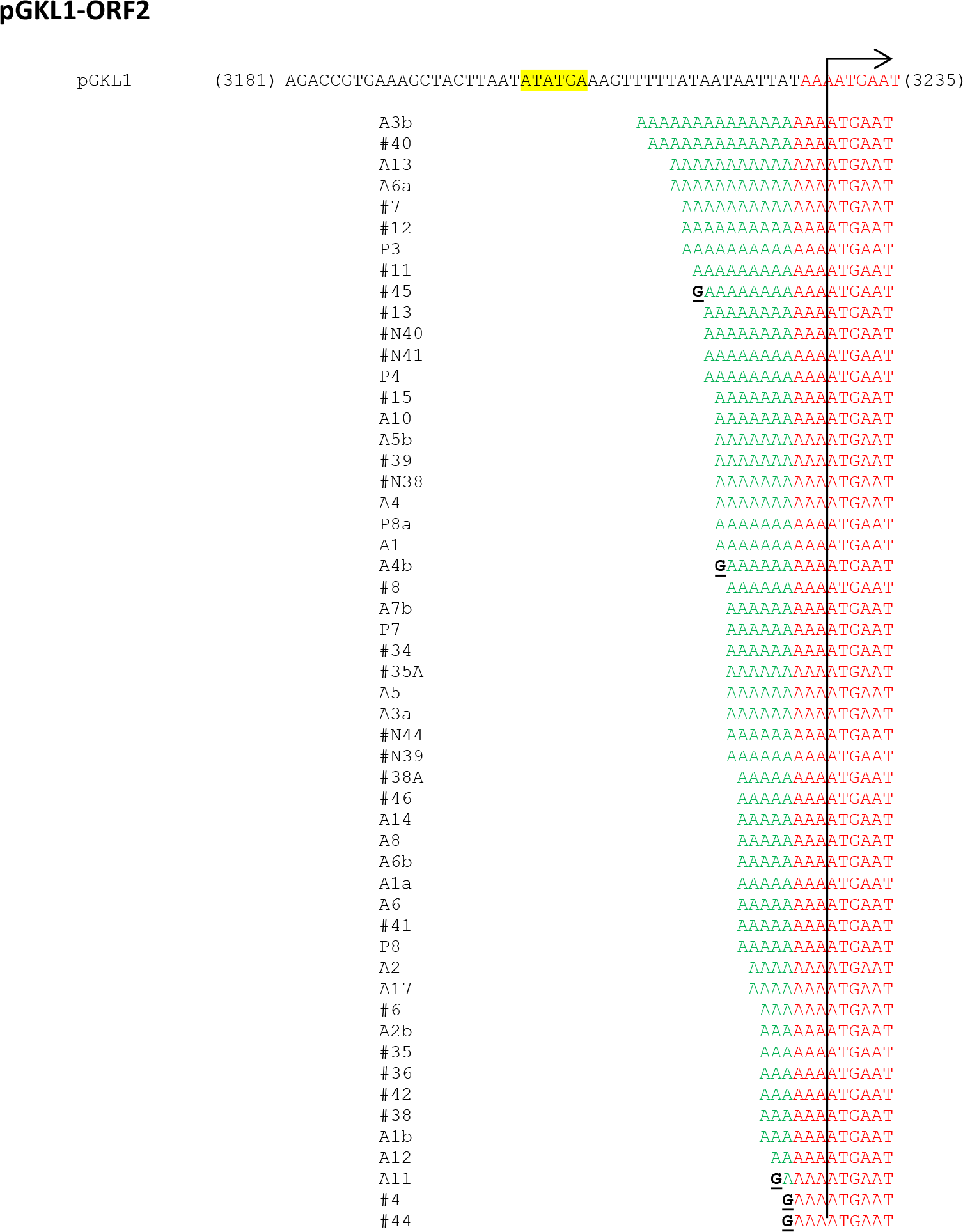
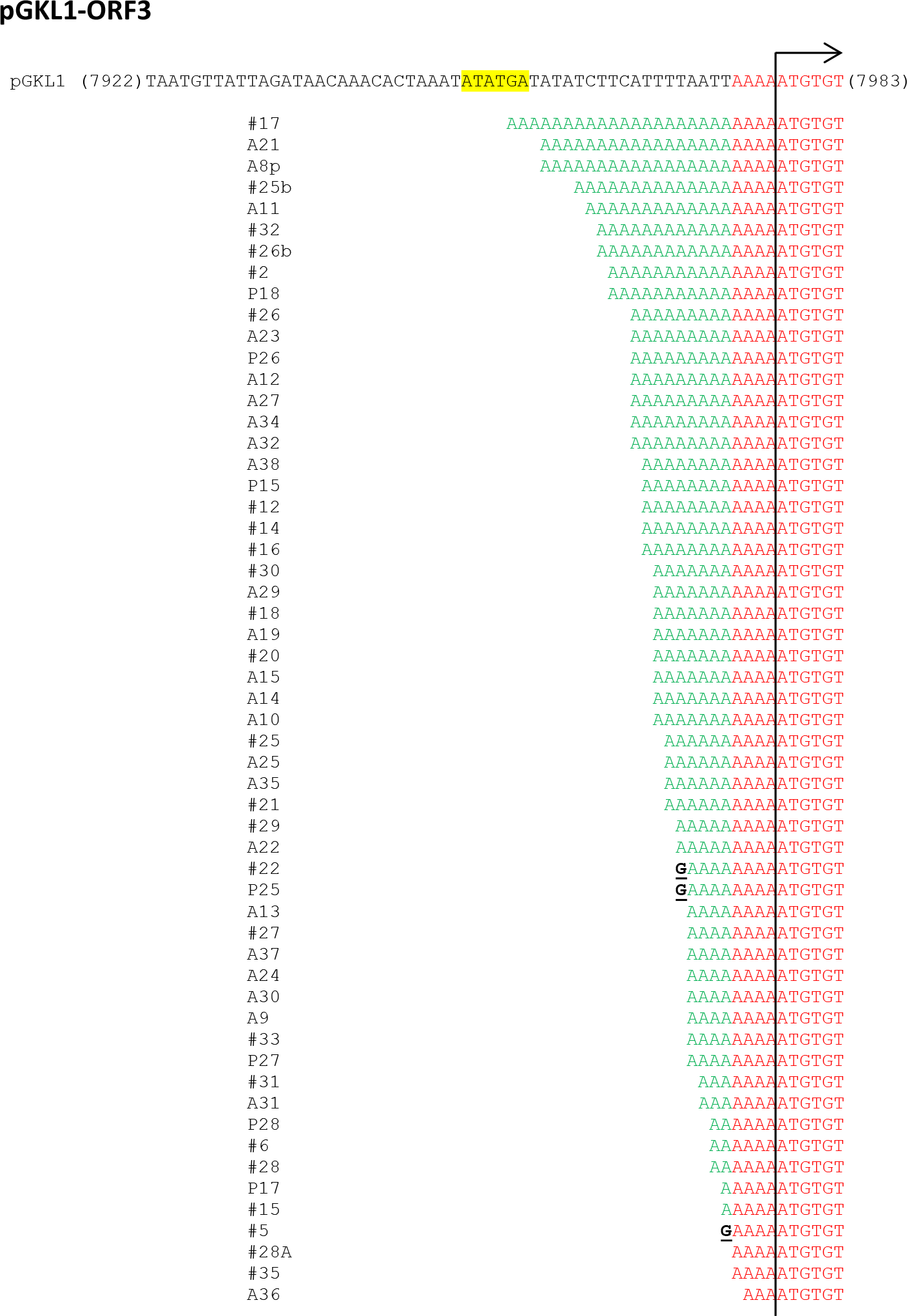
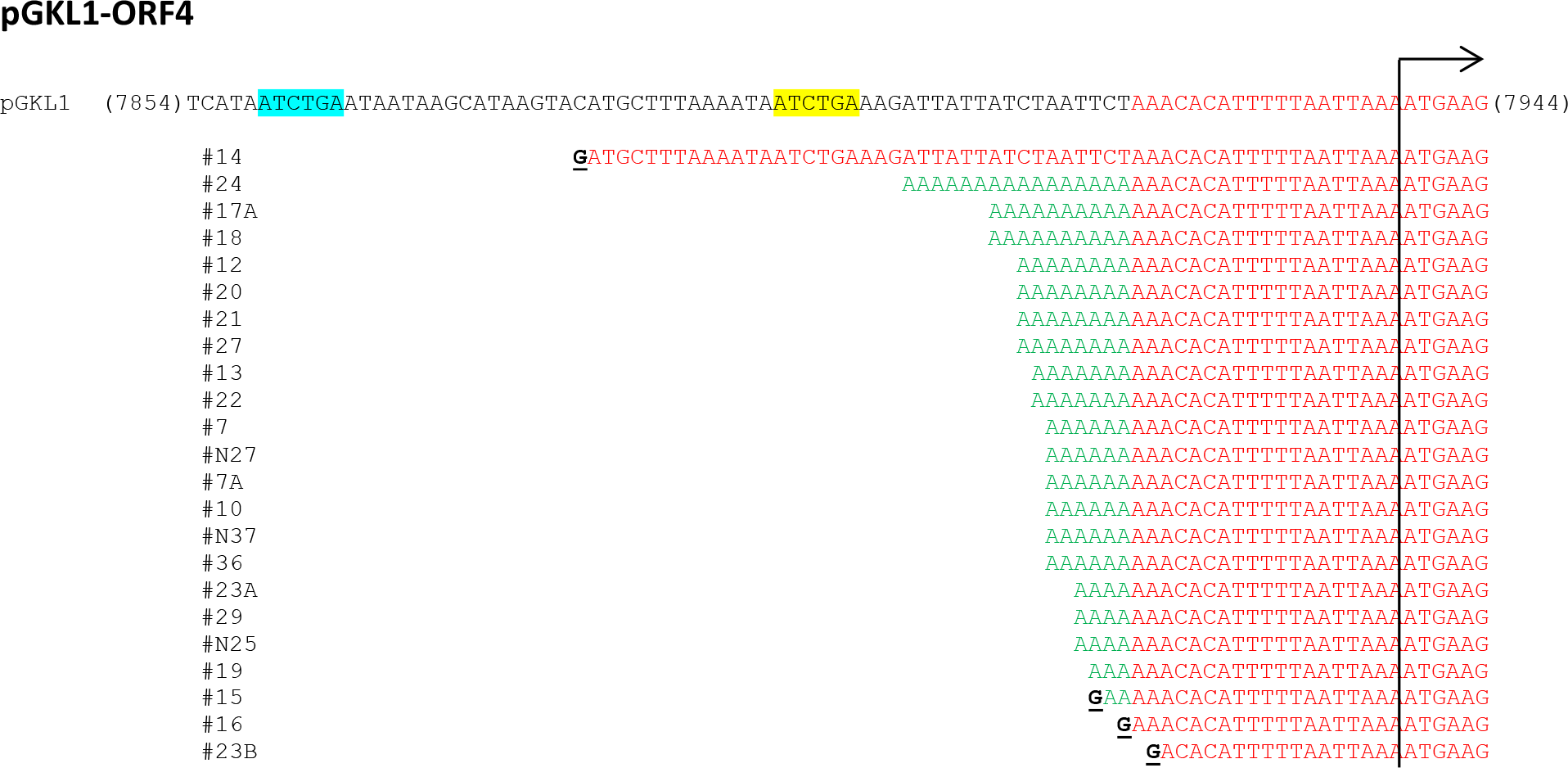
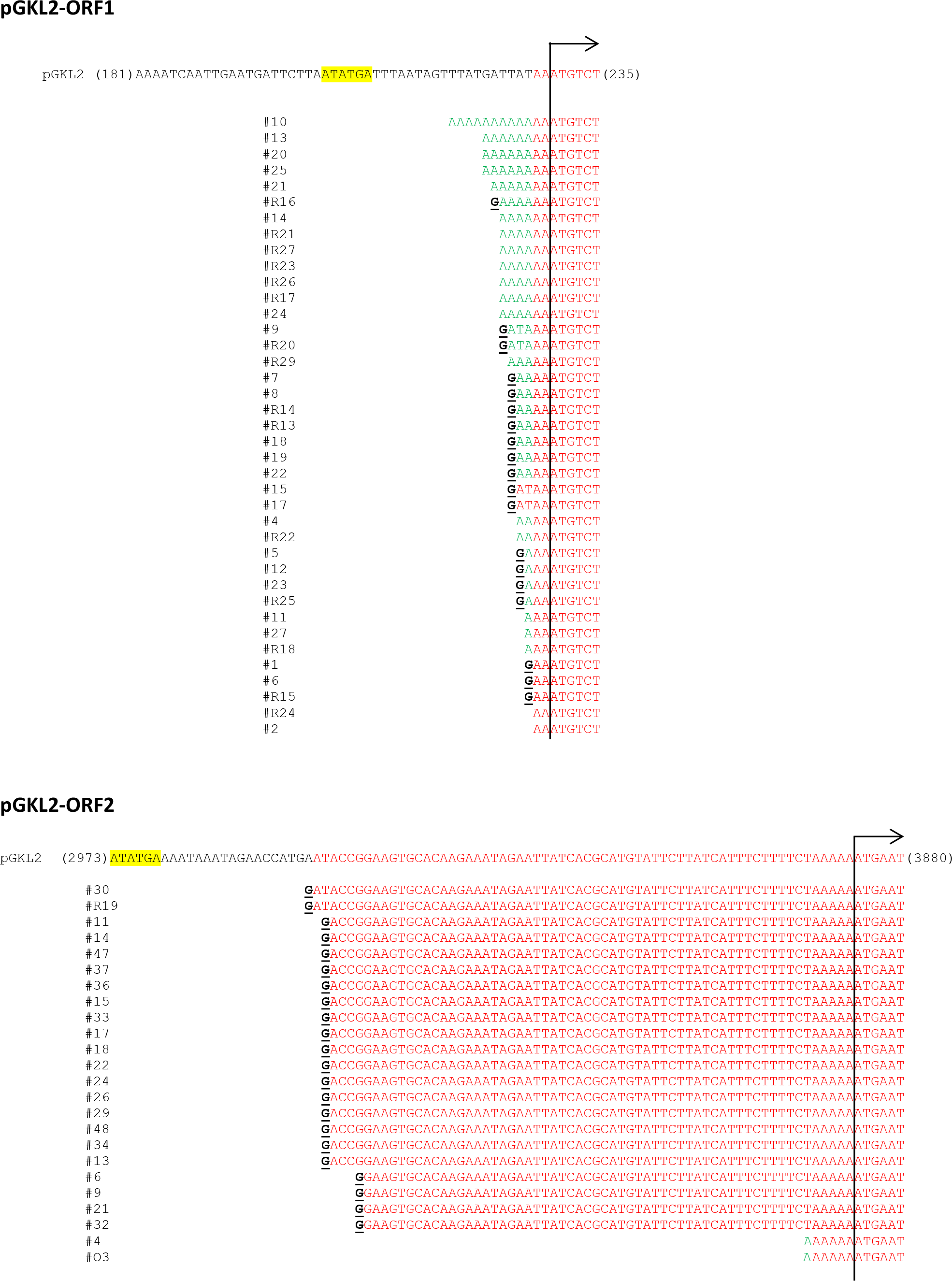
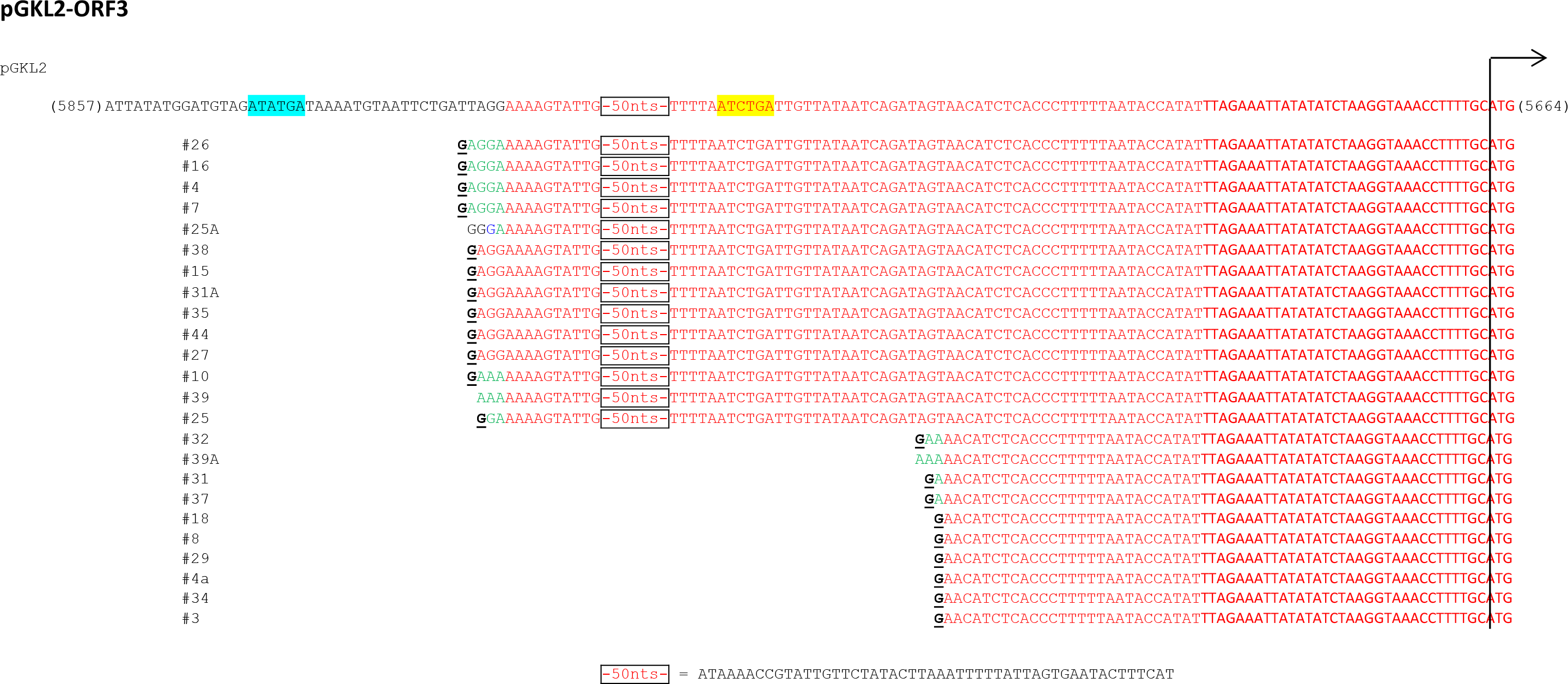
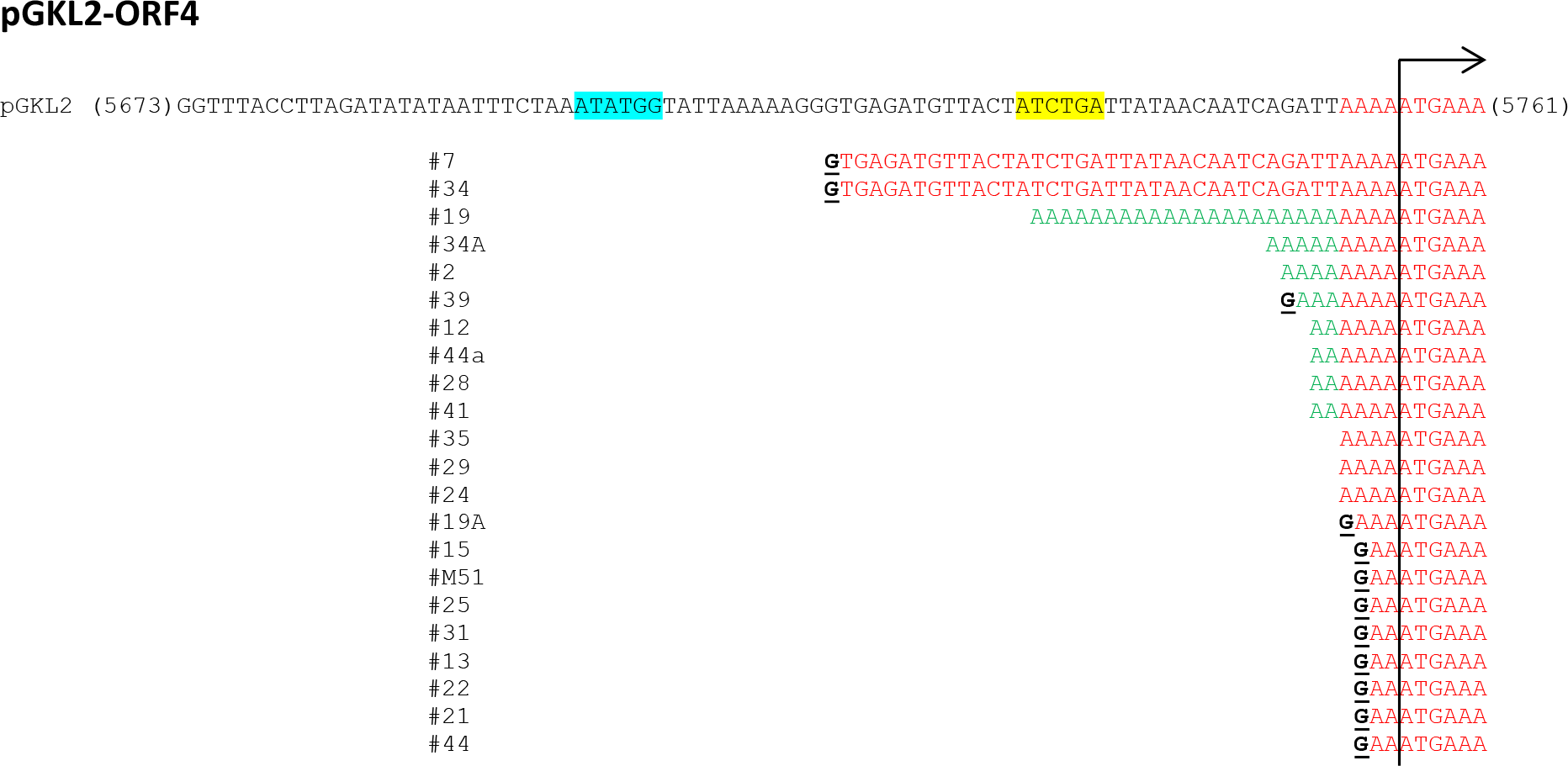
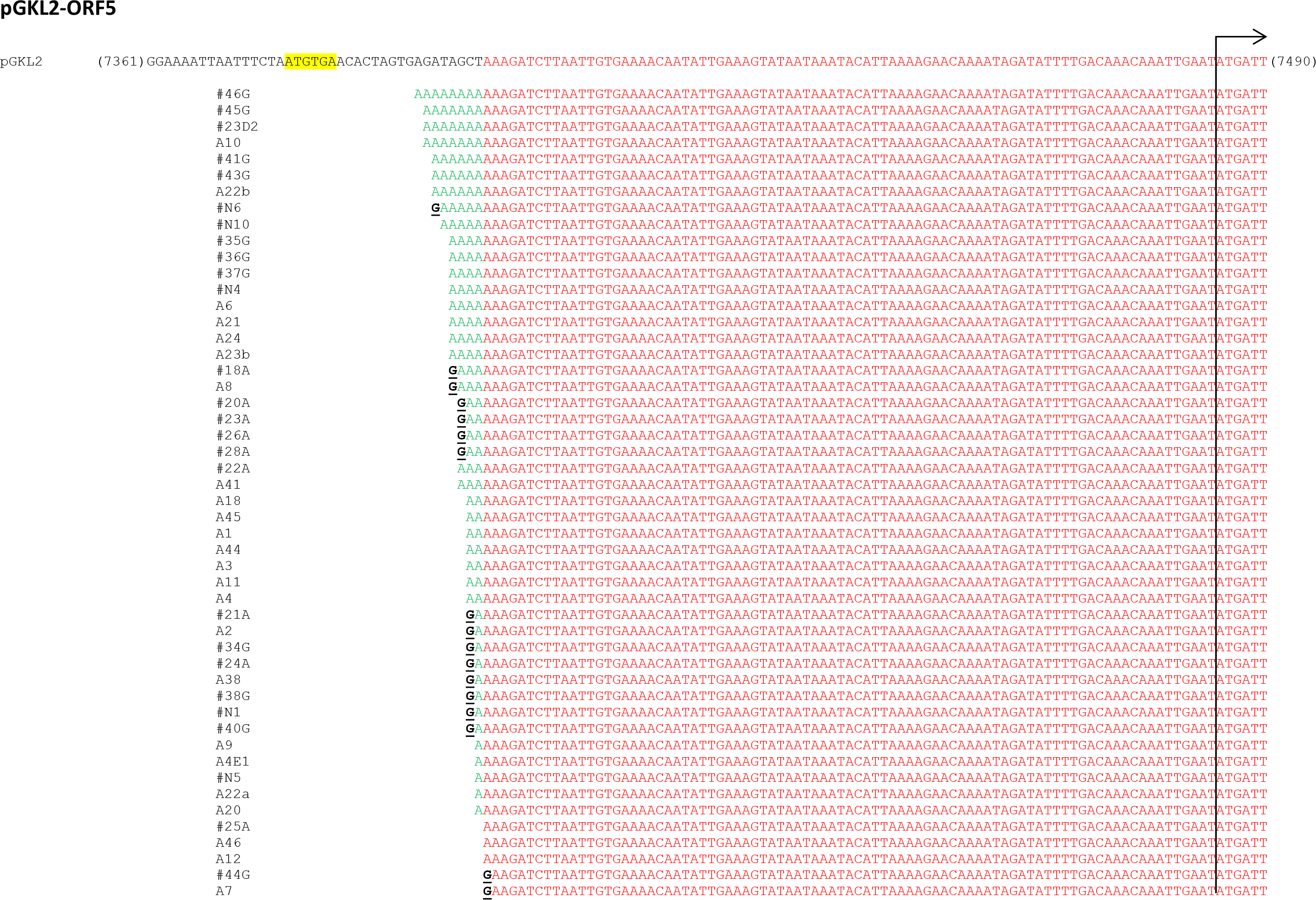
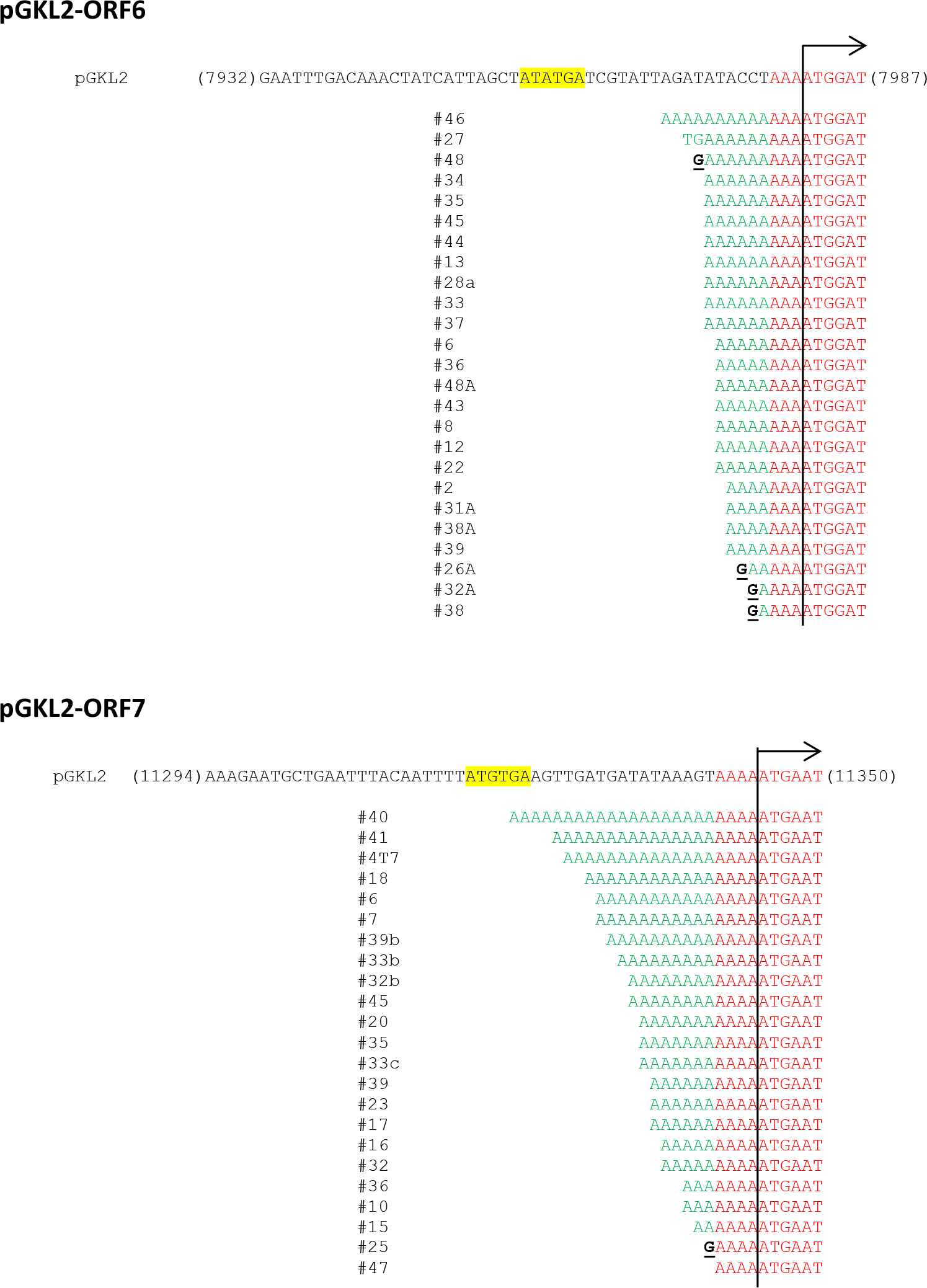
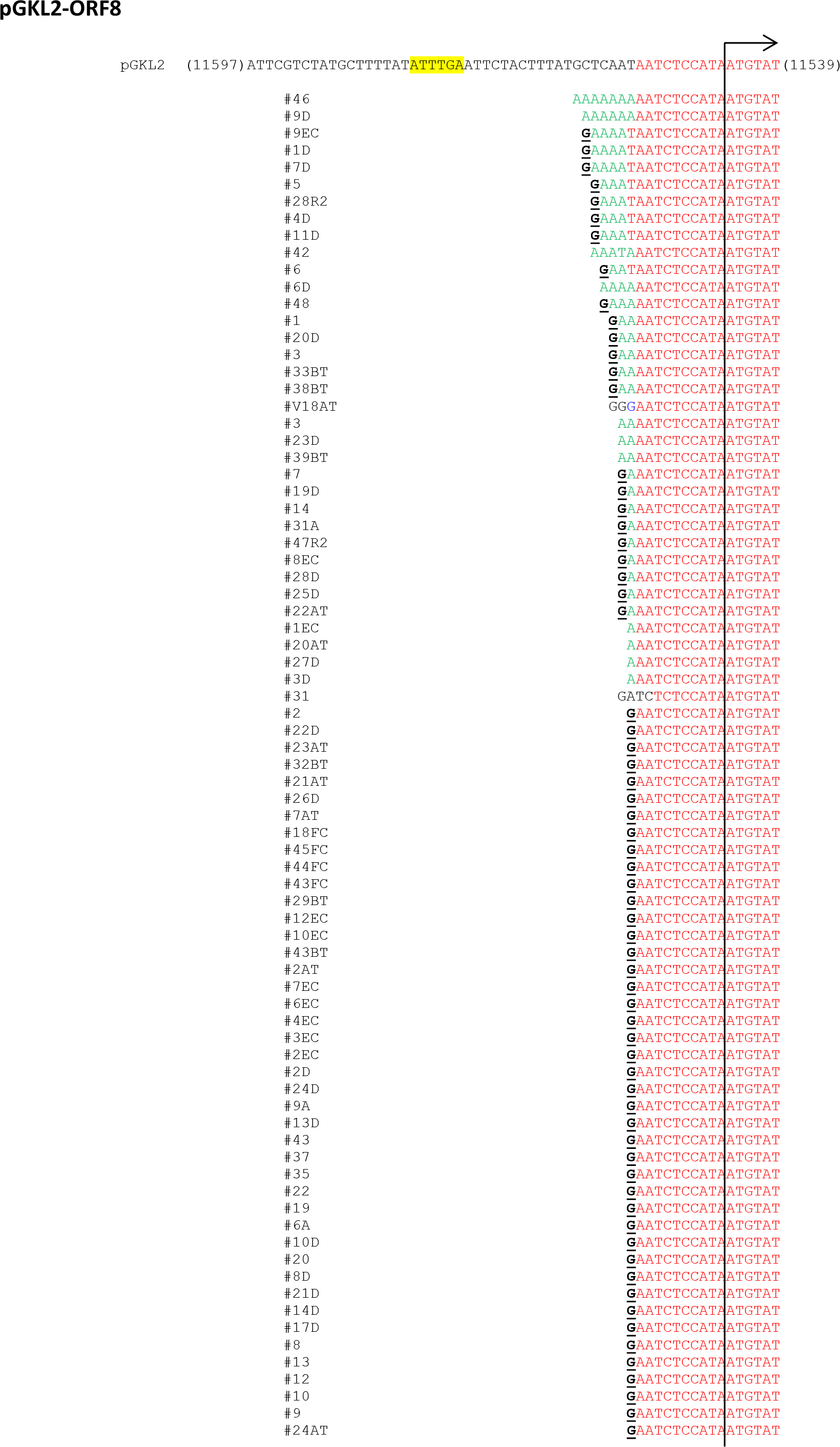
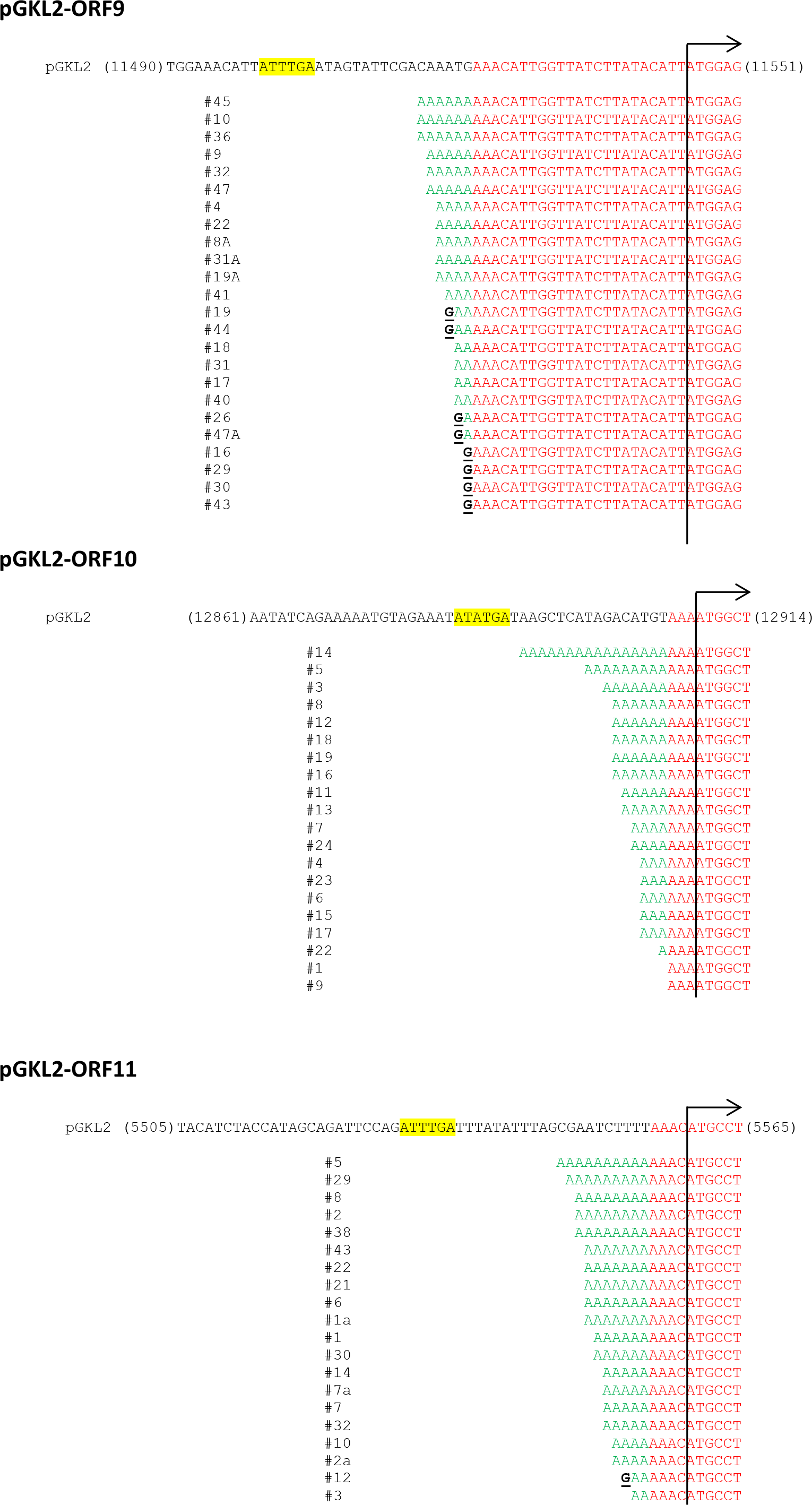
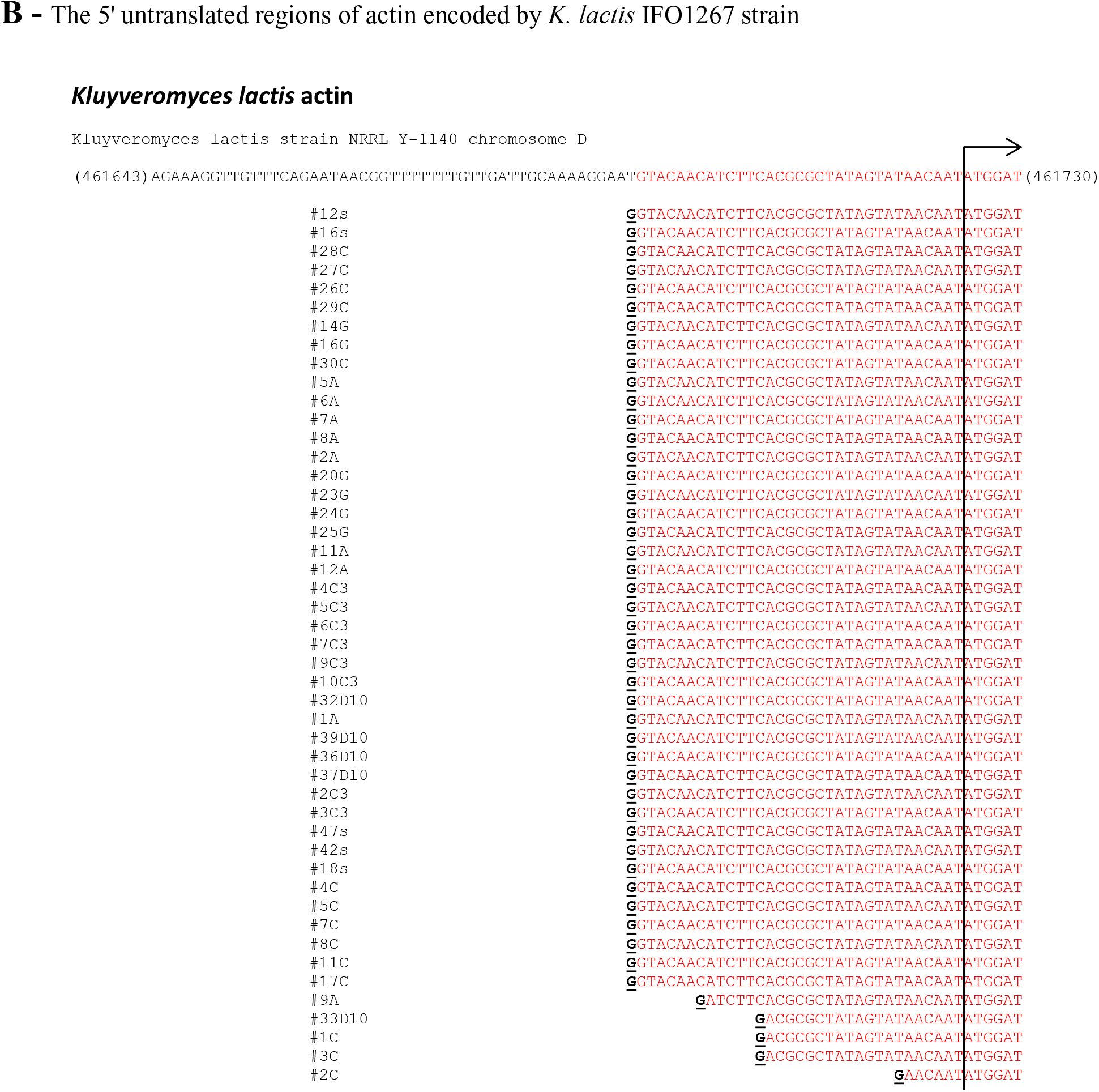
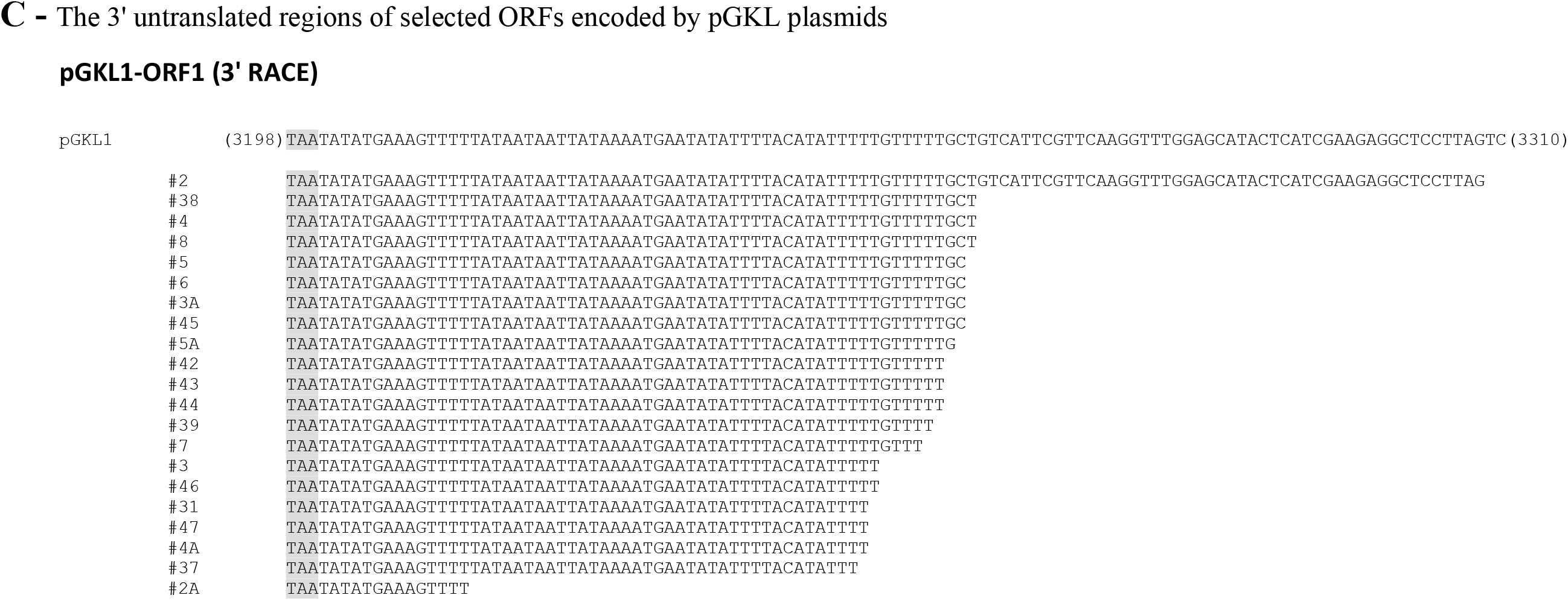
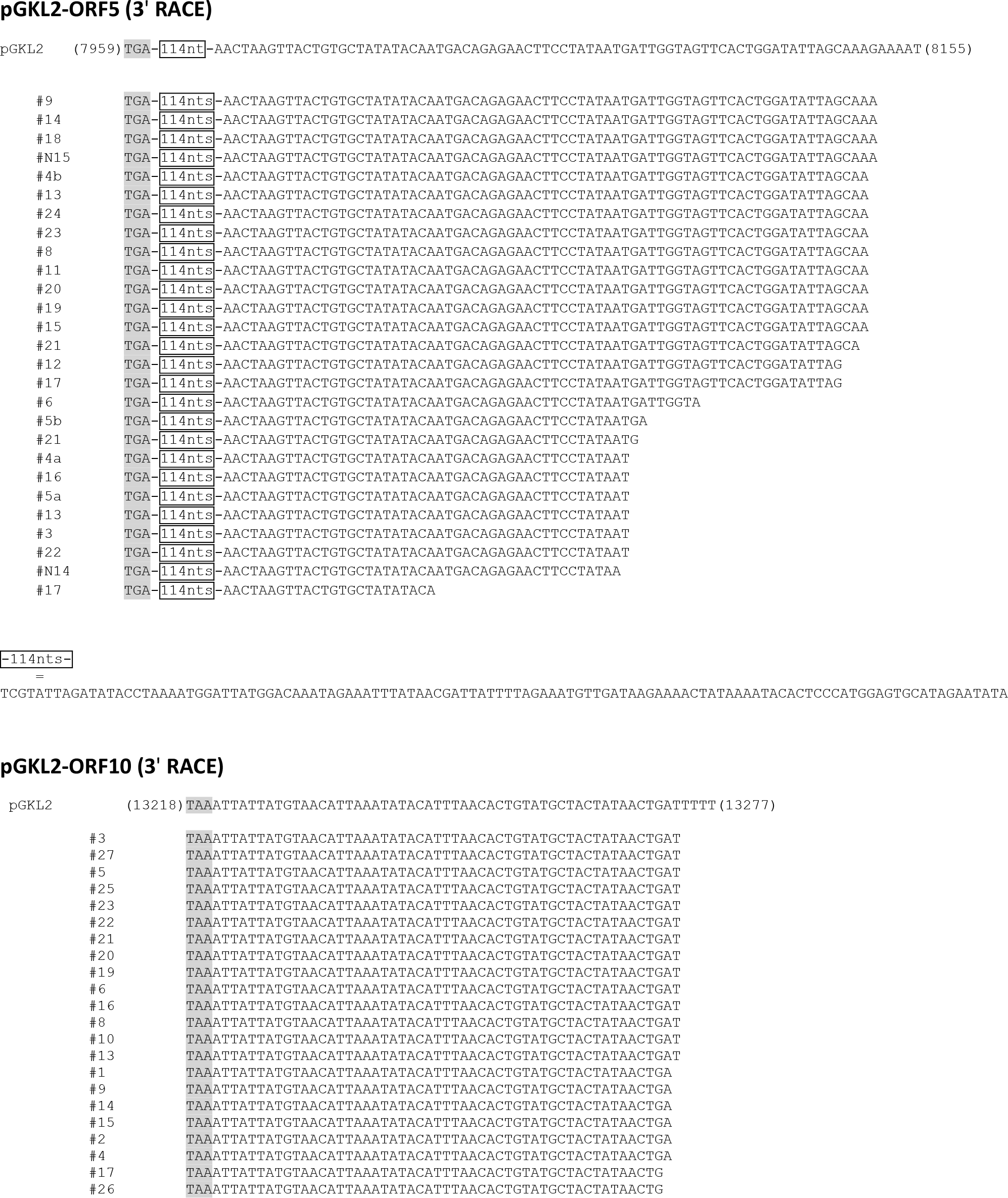
Molecular Analyses of pGKL mRNAs at the Level of Individual mRNA Molecules, Related to Figure 3. (A) The 5’ untranslated regions of ORFs encoded by pGKL plasmids. Plasmid DNA (pGKL1 or pGKL2) with an AUG initiation site (marked by a vertical line and an arrow that points in the direction of transcription) and part of the DNA upstream of the start codon, including the Upstream Conserved Sequence (UCS, in yellow box), is shown for each ORF. In the case of *K1ORF4*, *K2ORF3* and *K2ORF4*, new putative UCSs were found (shown in blue box) based on the detected transcripts. Numbers in parenthesis on both sides of the plasmid sequence indicate the position of the DNA sequence relative to GI: 163932456 (for pGKL1) or GI: 2868 (for pGKL2). Individual cDNA sequences corresponding to transcripts of a given ORF are aligned below the template DNA. After mRNA was isolated from *K. lactis* IFO1267, cDNA was prepared by SuperScript III Reverse Transcriptase (M-MuLV mutant; clone names begin with the number sign (#)) or by AMV Reverse Transcriptase (Finnzymes; clone names begin with the letter A; this enzyme was used only for *K1ORF2*, *K1ORF3* and *K2ORF5*). In this comparison, we have also included clones obtained from the double-mutant strain *K. lactis* IFO1267 *pbp1Δ pab1Δ* (see text; clone names beginning with the letter P; this approach was used only for *K1ORF2* and *K1ORF3*). The uniform distribution of the clones indicate that the reverse transcriptases and strains used for mRNA isolation/cDNA synthesis/clone preparation do not affect the results. The sequences of at least 20 independent clones were determined and aligned with the plasmid (template) DNA sequence of each ORF. Sequence sections that share sequence identity with the template plasmid DNA are labeled in red. Sequence sections that do not share sequence identity with the template plasmid DNA, thus containing non-template nucleotides of mRNA transcripts, are labeled in green. Guanosine residues that correspond to the original caps at the 5’ ends of the mRNAs are underlined in black (G). All sequences are shown in the 5’ to 3’ transcription orientation regardless of their ORF’s natural orientation in pGKL plasmids. (B) The 5’ untranslated regions of the actin gene (KLLA0_D05357g) encoded by *K. lactis* IFO1267. (C) The 3’ untranslated regions of *K1ORF1*, *K2ORF5* and *K2ORF10* encoded by pGKL plasmids. Template DNA with its stop codon (in the gray box) is shown for each ORF. Numbers in parenthesis present on both sides of the plasmid sequence indicate the position of the DNA sequence relative to the GenBank records indicated above. Unique obtained sequences (prepared using SuperScript III Reverse Transcriptase) corresponding to transcripts of a given ORF are aligned under the template DNA. The sequences of at least 20 independent clones were determined and aligned with the plasmid (template) DNA of each ORF.

(A) The 5’ untranslated regions of ORFs encoded by pGKL plasmids. Plasmid DNA (pGKL1 or pGKL2) with an AUG initiation site (marked by a vertical line and an arrow that points in the direction of transcription) and part of the DNA upstream of the start codon, including the Upstream Conserved Sequence (UCS, in yellow box), is shown for each ORF. In the case of *K1ORF4, K2ORF3* and *K2ORF4*, new putative UCSs were found (shown in blue box) based on the detected transcripts. Numbers in parenthesis on both sides of the plasmid sequence indicate the position of the DNA sequence relative to GI: 163932456 (for pGKL1) or GI: 2868(for pGKL2). Individual cDNA sequences corresponding to transcripts of a given ORF are aligned below the template DNA. After mRNA was isolated from *K. lactis* IFO1267, cDNA was prepared by SuperScript III Reverse Transcriptase (M-MuLV mutant; clone names begin with the number sign (#)) or by AMV Reverse Transcriptase (Finnzymes; clone names begin with the letter A; this enzyme was used only for *K1ORF2, K1ORF3* and *K2ORF5*). In this comparison, we have also included clones obtained from the double-mutant strain *K. lactis* IFO1267 *pbp1Δ pab1Δ* (see text; clone names beginning with the letter P; this approach was used only for *K1ORF2* and *K1ORF3*). The uniform distribution of the clones indicate that the reverse transcriptases and strains used for mRNA isolation/cDNA synthesis/clone preparation do not affect the results. The sequences of at least 20 independent clones were determined and aligned with the plasmid (template) DNA sequence of each ORF. Sequence sections that share sequence identity with the template plasmid DNA are labeled in red. Sequence sections that do not share sequence identity with the template plasmid DNA, thus containing non-template nucleotides of mRNA transcripts, are labeled in green. Guanosine residues that correspond to the original caps at the 5’ ends of the mRNAs are underlined in black (<u>G</u>). All sequences are shown in the 5’ to 3’ transcription orientation regardless of their ORF’s natural orientation in pGKL plasmids. (B) The 5’ untranslated regions of the actin gene (KLLA0_D05357g) encoded by *K. lactis* IFO1267. (C) The 3’ untranslated regions of *K1ORF1, K2ORF5* and *K2ORF10* encoded by pGKL plasmids. Template DNA with its stop codon (in the gray box) is shown for each ORF. Numbers in parenthesis present on both sides of the plasmid sequence indicate the position of the DNA sequence relative to the GenBank records indicated above. Unique obtained sequences (prepared using SuperScript III Reverse Transcriptase) corresponding to transcripts of a given ORF are aligned under the template DNA. The sequences of at least 20 independent clones were determined and aligned with the plasmid (template) DNA of each ORF.

## Supplemental Experimental Procedures

### Plasmid Construction

The *CDC33* (eIF4E) gene was amplified from *S. cerevisiae* genomic DNA using Pfu DNA polymerase (Fermentas) with eIF4Ef and eIF4Er primers containing NcoI and HindIII restriction sites as follows: 5 min at 95°C; then 25 cycles of 30 sec at 94°C, 30 sec at 55°C, and 2 min at 72°C; and finally, 10 min at 72°C. The PCR cassette was digested and inserted into the NcoI and HindIII restriction sites of pGEX-4T2 (GE Healthcare) to generate a pGEX4T2::CDC33 construct. A pYX212 yeast shuttle plasmid (Ingenius) was digested with EcoRI and XhoI restriction endonucleases. A sequence coding for K1 toxin was excised from a pYX213::M1 (Valis et al., 2006) plasmid using the same restriction enzymes and ligated into the digested pYX212 plasmid to generate the plasmid pYX212::M1. *K2ORF3* was amplified from *K. lactis* pGKL2 plasmid DNA using Pfu DNA polymerase (Fermentas) with the NLS-ORF3 forward and the klorf3_2 reverse primers as follows: 5 min at 95°C; then 5 cycles of 30 sec at 94°C, 30 sec at 55°C, and 2 min at 72°C; and finally, 10 min at 72°C. One microliter of this reaction mixture was used for subsequent PCR amplification using Pfu DNA polymerase with ORF3_NLS_BamF and the klorf3_2 reverse primers as follows: 5 min at 95°C; then 20 cycles of 30 sec at 94°C, 30 sec at 55°C, and 2 min at 72°C; and finally, 10 min at 72°C. The resulting PCR product was purified using agarose electrophoresis and cloned into a pCR4-TOPO plasmid (Invitrogen). An NLS-ORF3 cassette from a pCR4-TOPO plasmid was excised with BamHI and SalI restriction endonucleases and inserted into the corresponding sites of the pYX212 or pYX213 (Ingenius) plasmids to generate pYX212::NLS-ORF3 or pYX213::NLS-ORF3 constructs, respectively. All clones were verified by restriction endonuclease digestion and sequencing. All primers used in this study are listed in Table S2.

### Assay of Enzyme-GMP Complex Formation

A reaction mixture (20 μl) containing 50 mM Tris-HCl (pH 8.0); 2 mM DTT; 20 mM MgCl_2_; [α- ^32^P]GTP and purified K2Oof3p was incubated at 30°C for 60 min. The reaction was quenched with SDS-PAGE loading buffer, and the products were resolved by SDS-PAGE. The K2Orf3p-[^32^P]GMP adduct was visualized by autoradiography of the dried gel.

### Removal of Mitochondrial DNA by Ethidium Bromide Treatment

Isogenic *S. cerevisiae* strains bearing the wild-type *CDC33* gene (CWO4*CDC33*wt) and its temperature-sensitive mutations (CWO4*cdc33-1* and CWO4*cdc33-42*, for exact genotypes see Table S3) (Altmann et al., 1989; Altmann and Trachsel, 1989) were cultivated for two days in synthetic drop-out minimal medium (SD) that lacked leucine and tryptophan (SD-TL) and contained ethidium bromide at a final concentration of 25 μg per ml. Afterwards, these cultures were diluted five thousand times, cultivated again for two days and then diluted and cultivated once more. Loss of mitochondria was verified by the lack of colony growth on medium with glycerol as the sole carbon source and by DAPI staining followed by fluorescence microscopy.

### Transformation of Yeast Cells

All yeast transformations with plasmid DNA were performed with the one-step LiCl method (Gietz and Woods, 2002). The cells were grown in a shaker at 28°C in SD minimal medium lacking selected amino acids or nucleotide bases to ensure plasmid maintenance. Transformation using a PCR fragment for homologous recombination was carried out in the same way with the exception of including a five-hour incubation under nonselective conditions immediately after transformation prior to plating.

### Killer Production in Modified CWO4 Yeast Strains

Tested strains (50 ml) were cultured in an appropriate medium at 24°C with shaking. When the A_600_ reached 0.9, cells were harvested and washed three times with the same medium preheated to 24°C. One-half of the culture (25 ml) was mixed with 25 ml of fresh medium preheated to 24°C and then cultivated at 24°C for more than 20 hours. The second half of the tested culture (25 ml) was mixed with 25 ml of fresh medium preheated to 52°C and then cultivated at 37°C for more than 20 hours. Aliquots (2 ml) were taken at 0, 3, 6 and 12 hours. Samples were harvested, sterilized using 0.2 μm filters and assayed for the presence of the killer toxin activity in the culture medium by an agar well diffusion assay using *S. cerevisiae* S6/1 as a sensitive strain.

### Assay of killer toxin activity

Filter sterilized culture medium was tested for the presence of the killer toxin activity by the agar well diffusion assay using *S. cerevisiae* S6/1 as a sensitive strain. Approximately 2×10^5^ of sensitive yeast cells were plated onto OSS1 agar plates (5.3% worth agar; 0.7% agar; 1% glucose; 1M sorbitol; 0.002% methylene blue; buffered to pH 4.7 with sodium citrate) for testing of M1 killer toxin activity or onto YPD plates (1% yeast extract; 2% peptone; 2% glucose; 2% of agar) for testing of pGKL1 killer toxin activity. Wells were made with an 8 mm diameter cork borer and 100 μl of filter sterilized culture medium or 100 μl of serial dilution of the filter sterilized culture medium was pipetted into well. Inhibition zones were measured after 48 hours of incubation at 24°C.

### Statistical analysis

Results of at least five independent killer toxin activity measurements assayed by the well diffusion method were statistically evaluated using one-way ANOVA followed by post hoc Tukey’s HSD test and further confirmed with Scheffe multiple comparison. The normal distribution of the data was confirmed by Shapiro-Wilk test.

### Expression and Purification of mRNA Decapping Enzymes

pET28a::hDcp2 (Wang et al., 2002) and pET26b::Rai1 (Xiang et al., 2009b) (generous gifts from M. Kiledjian) expression plasmids were transformed into the *E.coli* Rosetta^™^ expression strain (Novagen). The corresponding mRNA decapping enzymes were expressed and purified according to previously published instructions (Piccirillo et al., 2003; Xiang et al., 2009b).

### *In Vitro* Processing of mRNAs by Decapping Enzymes

In a 100 μl reaction, 500 ng of total RNA purified from *K. lactis* IFO1267 or 4.1 pmol of *in vitro* transcribed and capped RNA were incubated with 0.25 μg of recombinant hDcp2 or 0.25 μg of recombinant Rai1 enzyme in 1x decapping buffer (10 mM Tris-HCl, pH 7.5; 100 mM potassium acetate; 2 mM magnesium acetate; 0.5 mM MnCl_2_; 2 mM dithiothreitol and 0.1 mM spermine) at 37°C for 30 min. After incubation, RNA was purified using an RNA Clean & Concentrator-5 kit (Zymo Research) and analyzed by 5’ RACE.

### *In Vitro* Synthesis of Capped mRNA

A DNA template corresponding to the first 579 nt (including 32 nt of 5’ UTR) of the *K. lactis ACT1* coding sequence (EnsemblFungi Id: KLLA0_D05357g) (Table S1B) was prepared by PCR with the forward primer modT7_KLactin_F (containing a modified T7 promoter sequence (Coleman et al., 2004)) and the reverse primer actin_KL-rev2 using *K. lactis* IFO1267 total cDNA as a PCR template as follows: 5 min at 95°C; then 30 cycles of 30 sec at 94°C, 30 sec at 55°C, and 1 min at 72°C; and finally, 10 min at 72°C. PCR products were purified using a High Pure PCR Product Purification Kit (Roche) and *in vitro*-transcribed using a TranscriptAid T7 High Yield Transcription Kit (Fermentas). Template DNA was removed by a DNA-free^™^ Kit (Ambion). Transcribed RNA was purified by an RNA Clean & Concentrator-5 kit (Zymo Research), and 9 pmol of RNA was capped *in vitro* using the vaccinia virus mRNA capping enzyme (Ambion) in the presence or absence of SAM according to the manufacturer’s instructions and purified using an RNA Clean & Concentrator-5 kit. Part of the sample was collected and analyzed by 5’ RACE to determine the efficiency of the mRNA capping reaction, while the rest of the sample was used for decapping assays with Rai1 or hDcp2 mRNA decapping enzymes.

### *PBP1, PAB1* and *LSM1* Gene Deletion

The *PBP1, PAB1* and *LSM1* genes were deleted from the chromosomes of the *K. lactis* IFO1267 strain using a loxP-G418-loxP cassette (Guldener et al., 1996). Briefly, the cassette for the deletion of the *PBP1* gene was amplified from the pUG6 plasmid by PCR with the primers KL_pbp1-del_Rev and KL_pbp1-del_For. The PCR product was resolved by agarose electrophoresis, purified, and transformed into *K. lactis* IFO1267. The deletion of the *PBP1* gene was verified by PCR. This modified strain was transformed with the pSH65 plasmid (Gueldener et al., 2002), incubated for 5 hours in nonselective medium and plated on solid medium supplemented with phleomycin (400 μg/ml). Monocolonies were grown on YPD plates with phleomycin for two days and subsequently tested for their ability to grow on G418 (250 μg/ml) selective plates. The excision of the G418 cassette was verified by PCR for selected colonies that did not grow in the presence of G418. These colonies were cultivated for five days with daily dilution under nonselective conditions (YPD medium only), resulting in the loss of the Cre-containing pSH65 plasmid. The resulting yeast strain, *K. lactis* IFO1267pbp1Δ, was used for subsequent *PAB1* deletion using the primers KL_pab1-del_For and KL_pab1-del_Rev in a similar fashion with the exception of the G418 cassette excision. *K. lactis* IFO1267 was used for *LSM1* deletion using primers KL_lsm1-del_For and KL_lsm1-del_Rev in a similar fashion with the exception of G418 cassette excision. The PCR reactions were carried out similarly (5 min at 95°C; then 30 cycles of 30 sec at 94°C, 30 sec at 55°C, and 4 min at 68°C; and finally, 10 min at 68°C). The nucleotide sequences of the primers used for the verification of the gene disruption cassettes are summarized in Table S2.

**Table S2.**
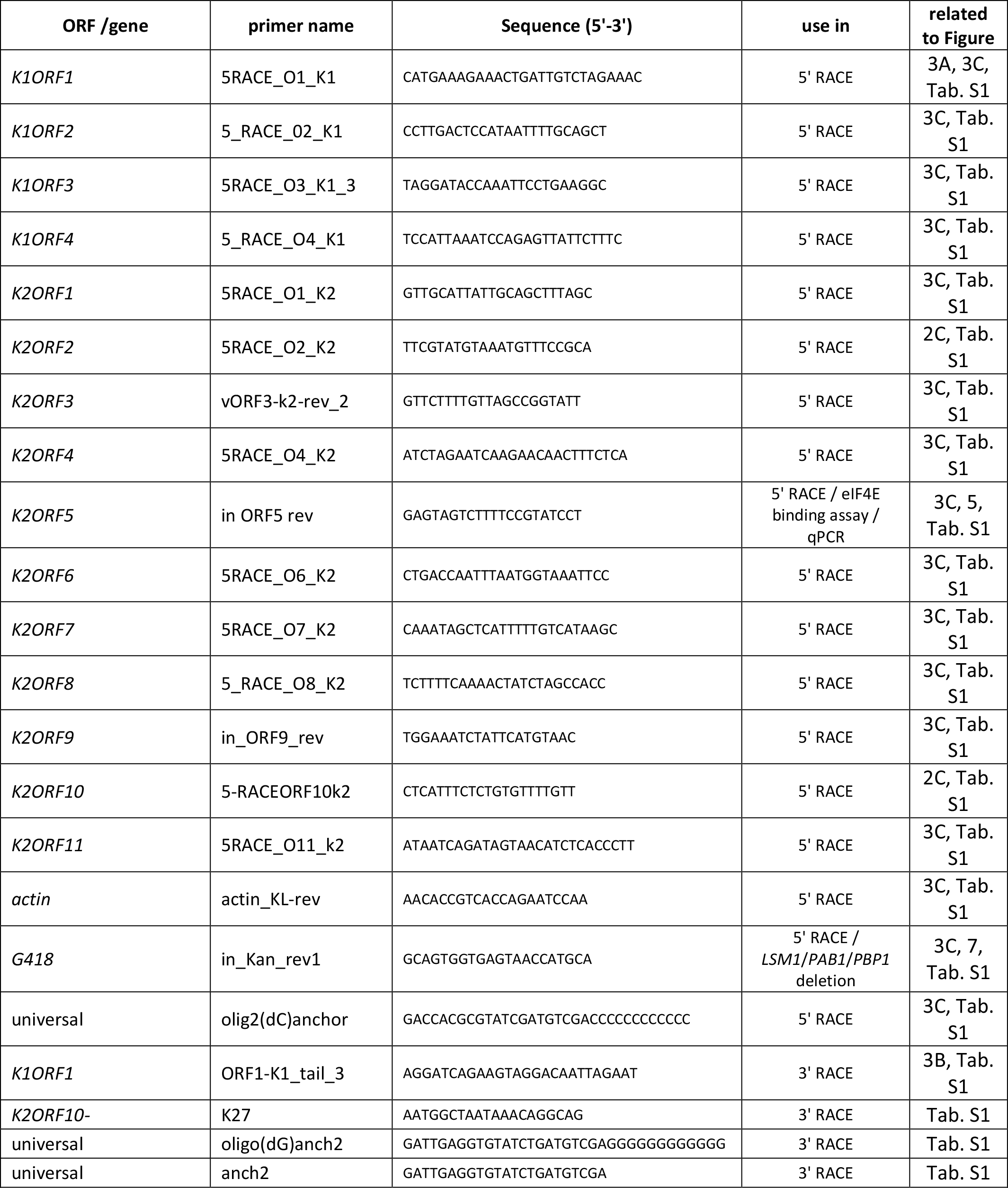
Primers Used in this Study, Related to Experimental Procedures

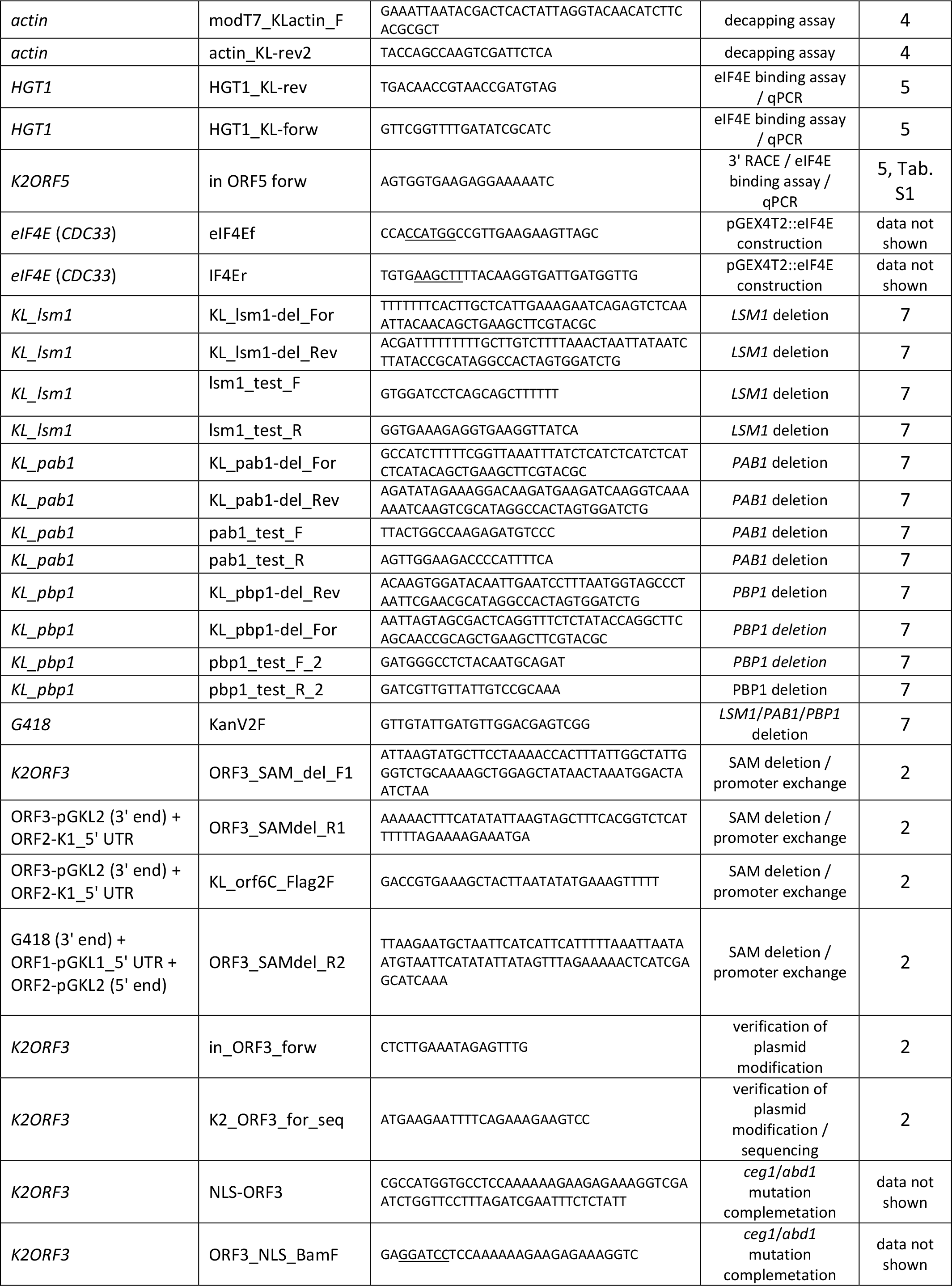

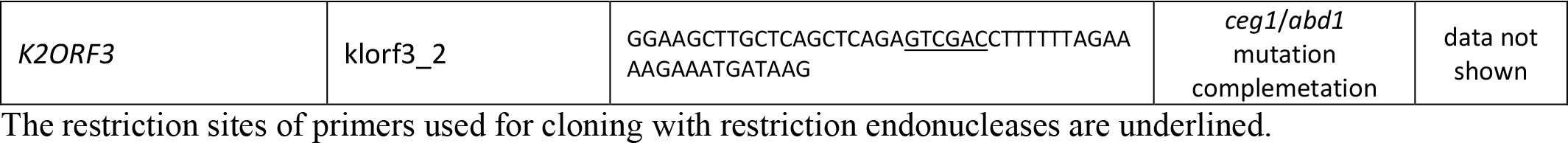

**Table S3.**
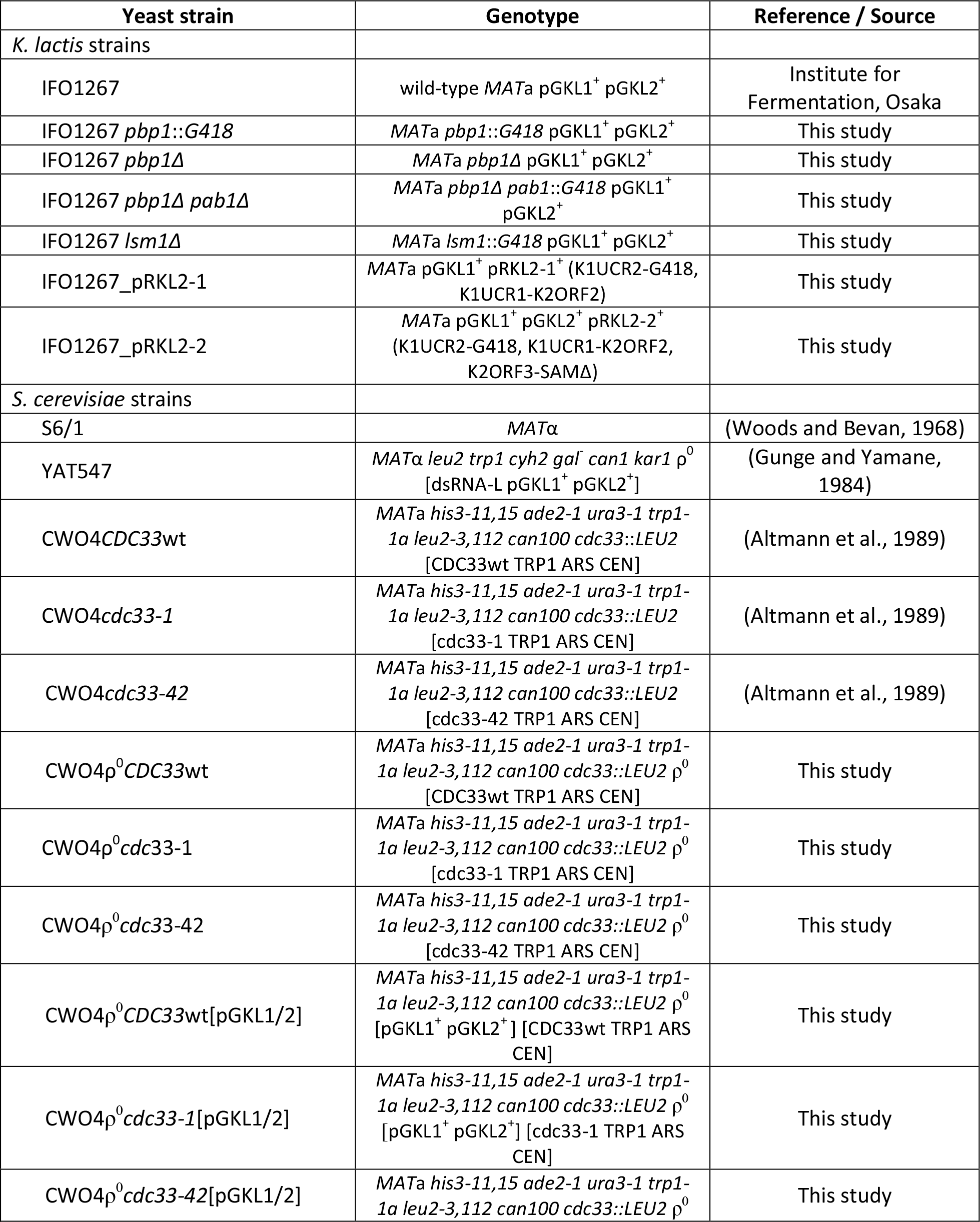
Yeast Strains Used in this Study, Related to Experimental Procedures

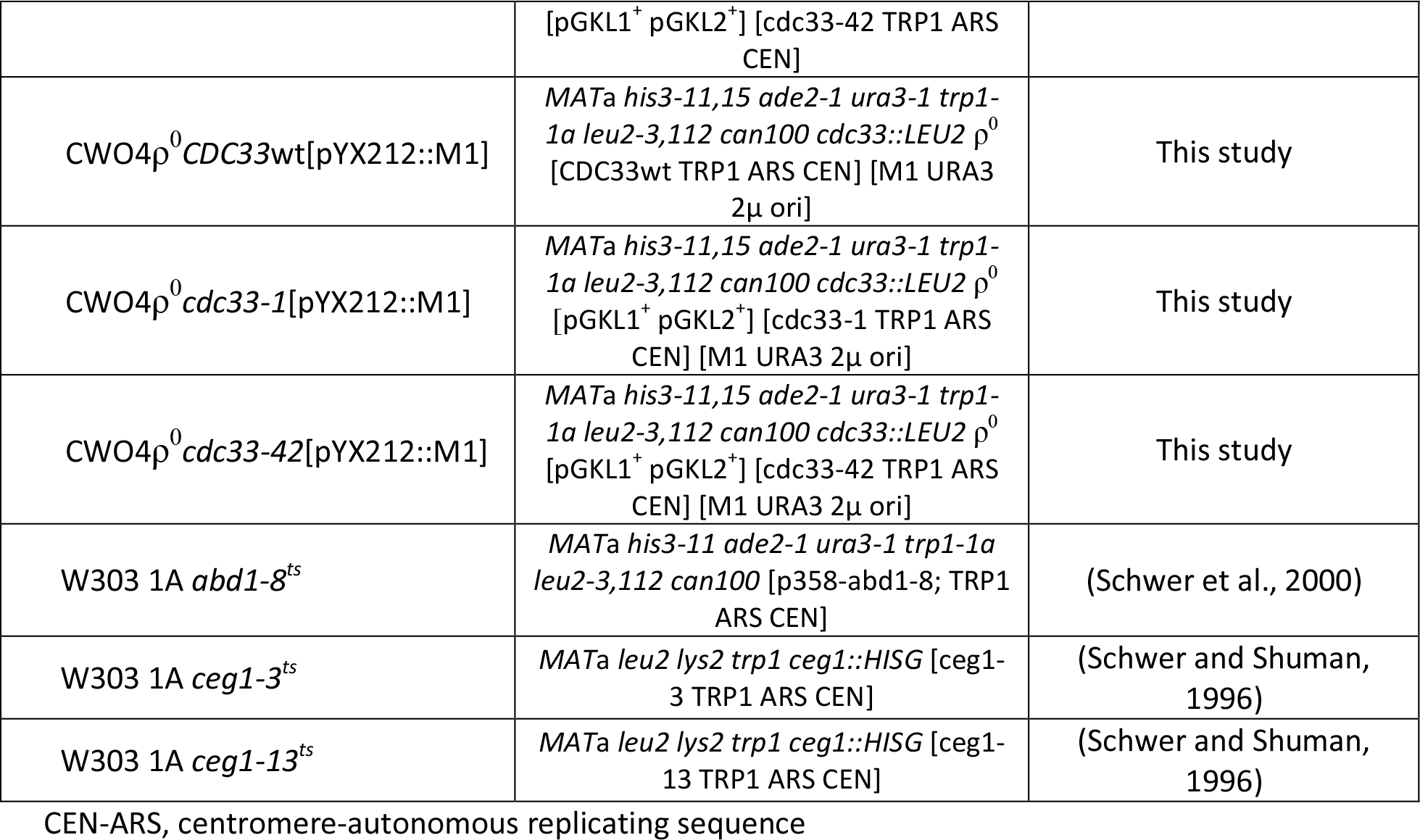

**Table S4.** Plasmids Used in this Study, Related to Experimental Procedures

| Plasmid | Description | Reference |
| --- | --- | --- |
| pET26b::Rai1 | aph(3')-II, pBR322 ori, f1 ori, lacI, Rop, pT7, Rai1-His | (Xiang et al., 2009a) |
| pET28a::hDcp2 | aph(3')-II, pBR322 ori, f1 ori, lacI, Rop, pT7, His-hDcp2 | (Wang et al., 2002) |
| pGKL1 | native, kil<sup>+</sup> | (Gunge et al., 1981) |
| pGKL2 | native | (Gunge et al., 1981) |
| pRKL2-1 | modified pGKL2 containing its own ORF2 under control of K1UCR1 and G418 resistance marker under control of K1UCR2 | This study |
| pRKL2-2 | same plasmid as above, additionally comprising D432A, G438A and D440A mutations in the SAM binding site located in K2ORF3p | This study |
| pUG6 | loxP-pAgTEF1-kanMX-tAgTEF1-loxP, G418 resistance, β-lactamase | (Guldener et al., 1996) |
| pSH65 | pGAL1-cre, ARS CEN, pAgTEF1-ble-tScCYC1, Phleo resistance, URA3, β-lactamase | (Gueldener et al., 2002) |
| pYX212 | 2μ ori, URA3, β-lactamase, TPI promoter | Ingenius Inc. |
| pYX213 | 2μ ori, URA3, β-lactamase, GAL1 promoter | Ingenius Inc. |
| pYX212::M1 | pYX212 containing cDNA copy of killer virus M1 | This study |
| pYX213::M1 | pYX213 containing cDNA copy of killer virus M1 | (Valis et al., 2006) |
| pYX212::NLS-K2ORF3 | pYX212 containing K2ORF3p fused with SV40 nuclear localization signal | This study |
| pYX213::NLS-K2ORF3 | pYX213 containing K2ORF3p fused with SV40 nuclear localization signal | This study |
| pCR4-TOPO | pLac, β-lactamase, aph(3')-II, ColE1, f1 ori | Invitrogen |
| pGEX-4T-2 | N-GST; pTac; lacI, thrombin; β-lactamase | GE Healthcare |
| pGEX4T2::CDC33 | pGEX-4T-2 containing yeast *S. cerevisiae* CDC33 gene produced as N-terminal GST fusion | This study |

UCR - sequence located between AUG initiation codon and UCS (including) of the selected ORF

**Table S5.** Analyses of the 5’ UTR poly(A) Leaders in pGKL mRNAs; Related to Figure 3.

| plasmid | ORF number | number of sequenced clones | number of clones with uncapped 5' poly(A) leader | % of clones with uncapped 5' poly(A) leader | number of adenosine residues<sup>1</sup> at 5' end of uncapped transcripts | clones with more than 12 adenosine residues at 5' end<sup>2</sup> |  | mean / median of adenosine residues<sup>1</sup> at 5' end, calculated per mRNA molecule for: |  |
| --- | --- | --- | --- | --- | --- | --- | --- | --- | --- |
|  |  |  |  |  |  | number | % | uncapped transcripts | all transcripts |
| pGKL1 | 1 | 27 | 21 | 77.7 | 6-15 | 6 | 22.2 | 10.19 / 10 | 8.81 / 10 |
|  | 2 | 53 | 48 | 90.5 | 6-18 | 13 | 24.5 | 10.38 / 10 | 10.05 / 10 |
|  | 3 | 56 | 53 | 94.6 | 4-25 | 29 | 51.8 | 10.38 / 12 | 11.66 / 12 |
|  | 4 | 23 | 19 | 82.6 | 6-19 | 3 | 13.0 | 10 / 9 | 8.69 / 9 |
| pGKL2 | 1 | 39 | 20 | 53.4 | 3-13 | 0 | 0 | 6.55 / 7 | 5.43 / 5 |
|  | 2 | 24 | 2 | 8.3 | 7 | 0 | 0 | N/A | 0.58 / 0 |
|  | 3 | 24 | 2 | 8.3 | 5-7 | 0 | 0 | N/A | 1 / 0 |
|  | 4 | 22 | 10 | 45.5 | 5-26 | 1 | 4.5 | 8.8 / 7 | 5.27 / 4.5 |
|  | 5 | 50 | 33 | 66 | 3-11 | 0 | 0 | 6.3 / 6 | 5.68 / 5 |
|  | 6 | 25 | 21 | 84 | 8-14 | 1 | 4.0 | 9.47 / 9 | 9 / 9 |
|  | 7 | 23 | 22 | 95.6 | 5-24 | 13 | 56.5 | 12.9 / 12 | 12.56 / 12 |
|  | 8 | 79 | 11 | 13.9 | 3-9 | 0 | 0 | 4.45 / 4 | 1.25 / 0 |
|  | 9 | 24 | 16 | 66.6 | 5-9 | 0 | 0 | 7 / 7 | 5.91 / 5.5 |
|  | 10 | 20 | 20 | 100 | 4-20 | 2 | 10 | 8.8 / 8.5 | 8.8 / 8.5 |
|  | 11 | 20 | 19 | 95 | 5-13 | 2 | 10 | 9.32 / 10 | 9.1 / 9.5 |

<sup>1</sup> template and non-template adenosine residues
<sup>2</sup> for capped and uncapped transcripts

